# N-hydroxy-pipecolic acid is a mobile signal that induces systemic disease resistance in *Arabidopsis*

**DOI:** 10.1101/288449

**Authors:** Yun Chu Chen, Eric C. Holmes, Jakub Rajniak, Jung-Gun Kim, Sandy Tang, Curt R. Fischer, Mary Beth Mudgett, Elizabeth S. Sattely

## Abstract

Systemic acquired resistance (SAR) is a global response in plants induced at the site of infection that leads to long-lasting and broad-spectrum disease resistance at distal, uninfected tissues. Despite the importance of this priming mechanism, the identity of the mobile defense signal that moves systemically throughout plants to initiate SAR has remained elusive. In this paper, we describe a new metabolite, *N*-hydroxy-pipecolic acid (N-OH-Pip), and provide evidence that this molecule is a mobile signal that plays a central role in initiating SAR signal transduction in *Arabidopsis thaliana*. We demonstrate that FLAVIN-DEPENDENT MONOOXYGENASE 1 (FMO1), a key regulator of SAR-associated defense priming, can synthesize N-OH-Pip from pipecolic acid *in planta,* and exogenously applied N-OH-PIP moves systemically in *Arabidopsis* and can rescue the SAR-deficiency of *fmo1* mutants. We also demonstrate that N-OH-Pip treatment causes systemic changes in the expression of pathogenesis-related genes and metabolic pathways throughout the plant, and enhances resistance to a bacterial pathogen. This work provides new insight into the chemical nature of a mobile signal for SAR and also suggests that the N-OH-Pip pathway is a promising target for metabolic engineering to enhance disease resistance.

## Introduction

Plants have developed a complex and dynamic innate immune system that relies on sensing and signaling using small molecules for defense against pathogens [1, 2]. At a primary site of infection, plants respond to common molecular features of microbes (e.g. bacterial flagellin or fungal chitin, collectively known as microbial associated molecular patterns or MAMPs) and pathogen-derived proteins, termed effectors, that enter the plant cell [3]. Immune responses to both classes of molecules (effector-triggered or pattern-triggered; [3, 4]) activate signaling transduction networks through the action of hormones such as salicylic acid (SA), ethylene, and jasmonic acid, which cause changes in defense gene expression and production of antimicrobial metabolites at the site of infection [5]. Interactions between these hormones function synergistically and antagonistically to tailor a specific immune response to different pathogens [2, 6].

While local defense is critical for limiting pathogen growth, plants also possess the ability to prime and amplify immune responses at distal sites. This global response is termed systemic acquired resistance (SAR) [2, 7]. SAR is critical for preventing the spread of pathogens and protecting against new infections [1, 2]. Although much is understood about how the immune response is activated locally, the chemical nature of the plant-derived molecules that mediate long distance communication between the site of infection (primary) and distal uninfected (secondary, systemic) sites have remain elusive.

Genetic and mechanistic studies of the model plant *Arabidopsis thaliana* have led to the identification of key components of the SAR pathway, providing critical insight into the mechanisms controlling long-lasting and broad-spectrum disease resistance [7]. Several plant-derived small molecules (e.g. SA [8], methyl salicylate (MeSA) [9], azelaic acid (AzA) [10], glycerol-3-phosphate (G3P) [11], dehydroabietinal (DA) [12], and pipecolic acid (Pip) [13] are associated with long-distance communication and signal amplification during SAR [8, 14, 15]. However, the onset of SAR signaling also requires the uncharacterized enzyme FLAVIN-DEPENDENT MONOOXYGENASE 1 (FMO1) [16–18]. Remarkably, treatment of *fmo1* mutants with AzA, DA, or Pip does not elicit systemic resistance, suggesting that a metabolite produced by FMO1 plays a key role in the establishment of SAR signaling [10, 12, 13]. The biochemical function of FMO1 has remained unknown.

Several forward genetic screens searching for SAR-deficient mutants identified multiple alleles of *fmo1, ald1* (*AGD2-LIKE DEFENSE RESPONSE PROTEIN 1),* and *sard4* (*SAR-DEFICIENT 4*), highlighting the importance of metabolites produced by these enzymes [16, 17, 19, 20]. ALD1 and SARD4 are involved in the biosynthesis of Pip [13, 21, 22]. Irrigation of wild type *Arabidopsis* plants with Pip induces SAR [13], which suggested that Pip might be a mobile SAR metabolite. Navarova et al. reported, however, that Pip could not trigger SAR in *fmo1* mutants [13]. In addition, *fmo1* plants accumulate high levels of Pip during a late stage of infection compared to wild-type plants [13]. These findings feature FMO1 as a key missing link in the mechanism of Pip-associated SAR.

We and others have found untargeted metabolite analysis of *Arabidopsis* genetic mutants to be a powerful approach for the identification of small molecules associated with fitness phenotypes (either previously characterized or suggested by transcriptome analysis) [23]. Examples include the identification and characterization of cytochromes P450 involved in phytoalexin production [24] and iron acquisition [25]. Given that FMO1 is one of the genes most responsive to biotic stress (as indicated by analysis of previously reported *Arabidopsis* microarray data summarized in Fig. S9), and that genetic data suggest molecules generated by FMO1 are required for initiating SAR [16–18], we sought to apply an untargeted metabolomics approach to determine the products of FMO1 and their function.

Here we report the discovery of glycosylated N-hydroxypipecolic acid (N-OGlc-Pip) in *Arabidopsis*. We provide evidence that the aglycone, N-OH-Pip, is the direct product of FMO1 and has a central role in SAR signal transduction [26]. We also demonstrate that exogenously applied N-OH-Pip can rescue the SAR deficient response of *fmo1* mutants, cause systemic changes in expression of pathogenesis-related genes and metabolic pathways throughout the plant (and distal to the site of application), and enhance resistance to the bacterial pathogen, *Pseudomonas syringae*. In addition, we provide biochemical evidence that *Arabidopsis* FMO1 catalyzes N-hydroxylation of the non-proteinogenic amino acid Pip to N-OH-Pip using transient expression assays in *Nicotiana benthamiana.* Taken together, our data indicates that N-OH-Pip is a key signaling molecule that is required to initiate SAR signaling in *Arabidopsis*.

## Results

### Untargeted metabolomics of *Arabidopsis fmo1* seedlings

To discover metabolites whose production is dependent on FMO1, we used liquid chromatography-mass spectrometry (LC-MS) based untargeted metabolomics to compare the composition of methanolic extracts from two sets of 12-day old seedlings grown hydroponically: *A. thaliana* Col-0 WT (WT) and a T-DNA insertion mutant of *FMO1*, *fmo1-1* [16], herein referred to as *fmo1*. The bacterial pathogen *Pseudomonas syringae* pathovar *tomato* DC3000 (*Pst*) was added to the seedling media 24 hours prior to metabolite analysis in order to elicit expression of enzymes associated with defense and SAR (including FMO1). This analysis revealed a major mass signal that is present in WT plants in response to *Pst* treatment but not Mock treatment (10 mM MgCl_2_) and was absent from all *fmo1* plants (Fig. 1A; Fig. S1B). We propose this signal corresponds to the O-glycosylated form of the novel metabolite N-OH-Pip, based on comparison of the MS/MS spectrum to an authentic synthetic standard of the N-OH-Pip aglycone (Fig. S2A-B, S3, S4). Although N-OH-Pip has not previously been observed as a naturally occurring metabolite in plants or other organisms, pipecolic acid has been observed in *Arabidopsis* and tomato [13, 27] and was shown to be associated with the SAR response [13]. Therefore, we hypothesized that FMO1 catalyzes the N-hydroxylation of Pip to produce N-OH-Pip. Glycosylated N-OH-Pip (N-OGlc-Pip) detectable in *Arabidopsis* plant extracts is likely the product of unknown UDP-glycosyltransferases that further processes N-OH-Pip (Fig. 1C). We did not detect free N-OH-Pip in WT seedlings elicited with *Pst* by LC-MS analysis (Fig. S1B). These data suggest that the N-OH-Pip aglycone does not accumulate in cells and/or is an unstable metabolite.

**Figure 1.**
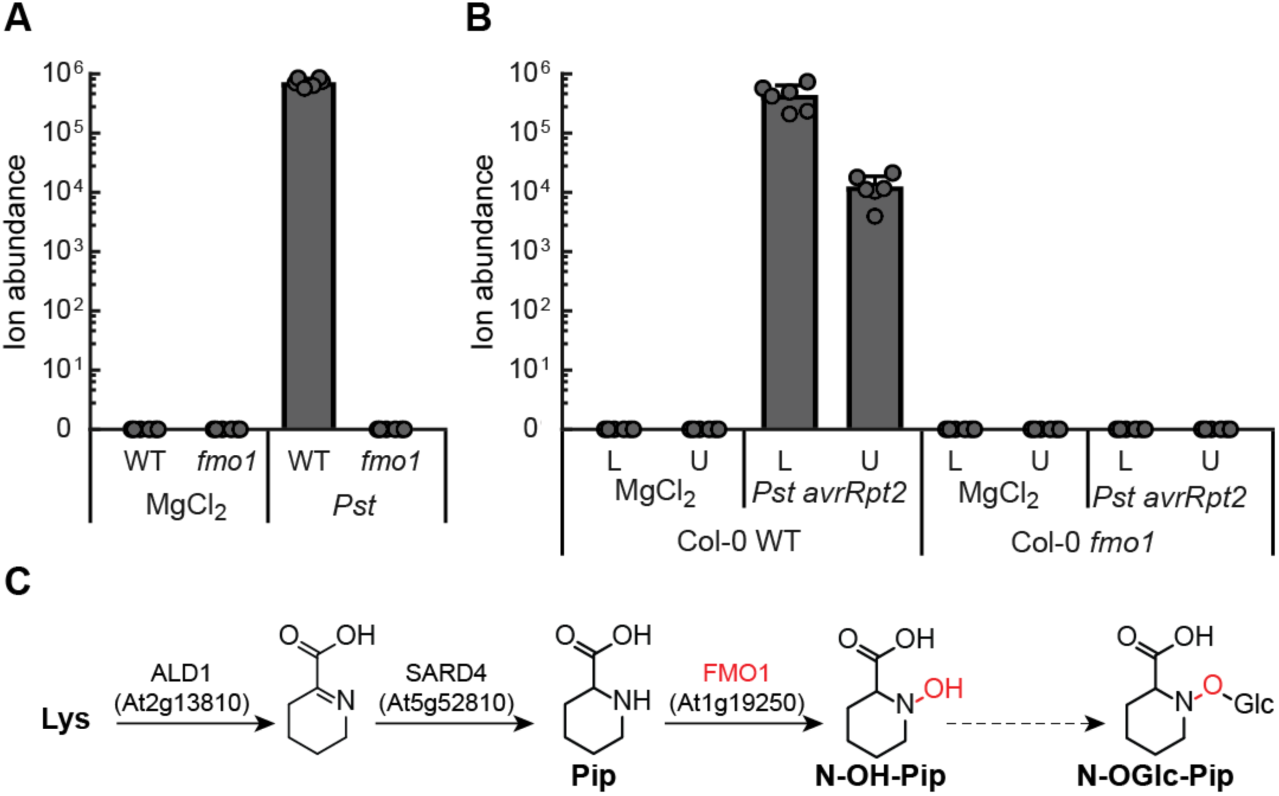
Untargeted metabolomics of *Arabidopsis* seedlings elicited with *Pst* implicates FMO1 in the production of pipecolic acid derivatives. (A) Ion abundances for N-OGlc-Pip (gray bars) detected in extracts isolated from *Arabidopsis* Col-0 WT and *fmo1* seedlings grown hydroponically and elicited with *Pseudomonas syringae* pv. *tomato* DC3000 (*Pst*). Levels represent the mean +/- STD of six biological replicates. Levels reported as zero indicate no detection of metabolites. (B) Ion abundances for N-OGlc-Pip (black bars) in lower and upper leaves of adult *Arabidopsis* Col-0 plants 48 hours post-inoculation (48 hpi) with a 5 x 10^6^ cfu/ml suspension of *Pst* expressing the T3S effector *avrRpt2* (*Pst avrRpt2*). Levels represent the mean +/- STD of six biological replicates. Levels reported as zero indicate no detection of metabolites. (C) Proposed biosynthetic activity for *Arabidopsis* FMO1. The *Arabidopsis* enzymes ALD1 and SARD4 convert lysine to pipecolic acid (Pip) (Ding et al., 2016). FMO1 is proposed to hydroxylate Pip to N-hydroxypipecolic acid (N-OH-Pip). Unknown enzymes are proposed to glycosylate N-OH-Pip to produce N-OGlc-Pip, the novel molecule identified in the untargeted metabolite analysis.

### Measurement of N-OH-Pip derivatives in adult plants

We also measured the level of N-OGlc-Pip in adult WT and *fmo1* plants (both in lower and upper leaves), after treatment of lower leaves with 10 mM MgCl_2_ (Mock) or *Pst* expressing the type III secretion (T3S) effector gene *avrRpt2* (*Pst avrRpt2*). *Pst avrRpt2* is an avirulent strain that induces effector-triggered immunity (ETI) and SAR signaling in resistant RPS2 *Arabidopsis* plants [28–30]. N-OGlc-Pip was detected in the lower and upper leaves of WT only after infection of lower leaves with *Pst avrRpt2*, but not in *fmo1* plants (Fig. 1B). As in seedlings, we were not able to detect free N-OH-Pip by LC-MS. Collectively, these data indicate that FMO1 is required for the production of N-OGlc-Pip in *Arabidopsis*.

### Biochemical activity of *Arabidopsis* FMO1

*Arabidopsis* FMO1 expressed and purified as a 6x-His fusion protein (FMO1-6x-His) from *E. coli* lacked the co-factor FAD and was not catalytically active in any of our *in vitro* biochemical assays. However, we were able to detect FMO1-dependent activity when *Arabidopsis* FMO1 was transiently expressed in *Nicotiana benthamiana* after feeding pipecolic acid and then analyzing products present in leaf methanolic extracts (Fig. 2, Fig. S1D). As shown in Figure 2, FMO1-expressing leaves accumulated free N-OH-Pip. They also contained a second metabolite with an m/z of 100 and a putative structure of N-hydroxy piperidene (Fig. 2A, Fig. S2C), which we propose is the result of oxidative decarboxylation of N-OH-Pip. We noted that the mass signal m/z 100 can be produced in samples of synthetic N-OH-Pip after heating in buffer. It is also present in Col-0 seedlings after *Pst* treatment and absent from *fmo1* seedlings (Fig. S1B), suggesting that the m/z 100 metabolite is likely FMO1-derived. In addition, N-OH-Pip levels in *N. benthamiana* extracts decreased between 28 and 48 hours post-infiltration (hpi) while the abundance of m/z 100 increased (Fig. S1C), further suggesting that N-OH-Pip is unstable *in planta* and may convert to m/z 100 over time. *N. benthamiana* leaves expressing two FMO1 active-site mutants, FMO1 G17A/G19A or FMO1 G215A, did not produce N-OH-Pip or m/z 100 (Fig. 2A). Taken together, these data demonstrate that FMO1 can catalyze the hydroxylation of Pip *in planta* (Fig. 2) and support the requirement of the putative FAD and NADP^+^ domains for FMO1 catalytic activity [18]. N-OGlc-Pip was not detected in *N. benthamiana* leaf extracts when *Arabidopsis* FMO1 was expressed, suggesting that *N. benthamiana* either does not have the necessary glycosyl transferases or that they are not expressed under our experimental conditions.

**Figure 2.**
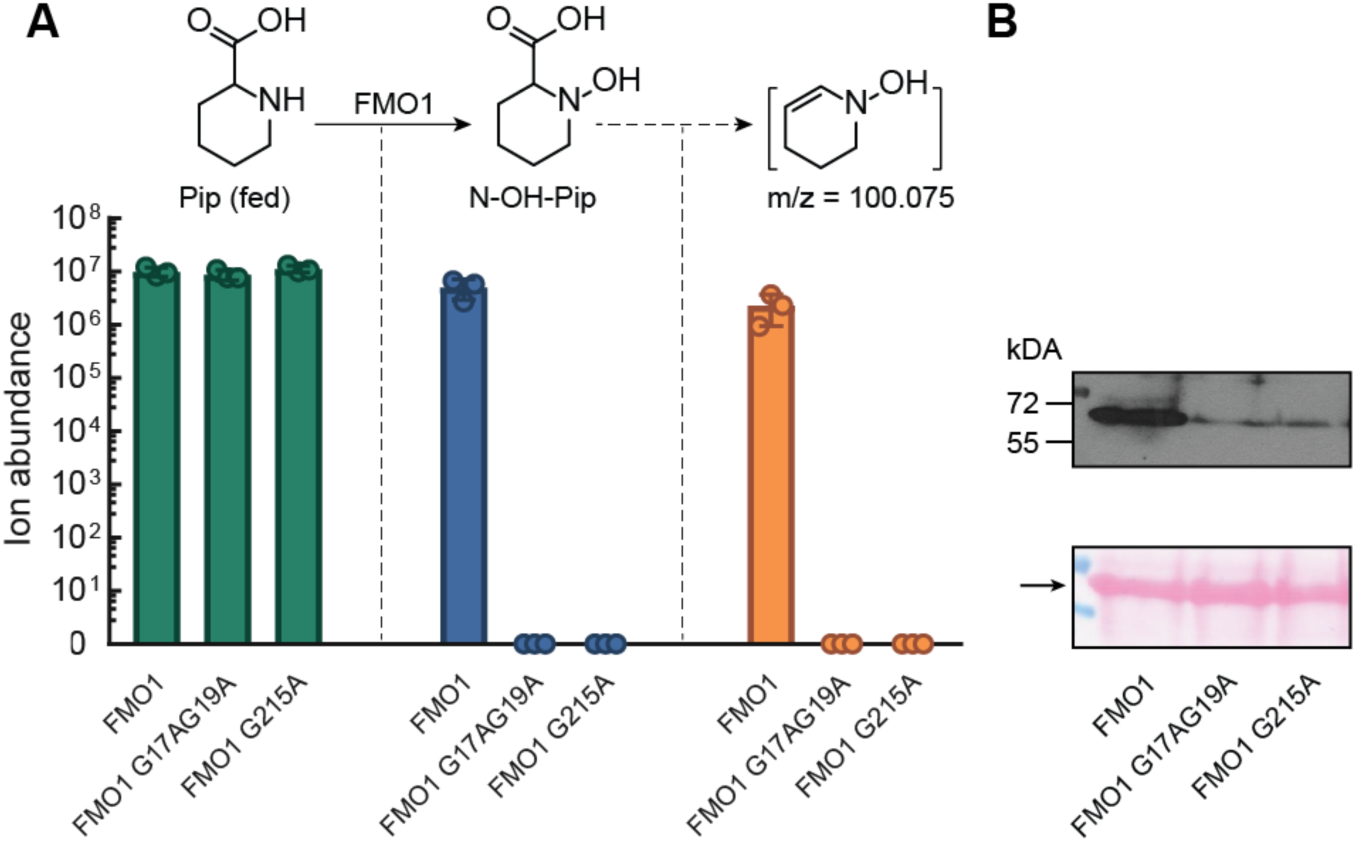
Overexpression of *Arabidopsis* FMO1 in *N. benthamiana* converts Pip to N-OH-Pip. (A) Ion abundance of Pip (green bars), N-OH-Pip (blue bars), and an unknown metabolite, m/z = 100.075 (orange bars) in leaves of *N. benthamiana* expressing *Arabidopsis* Col-0 WT FMO1-6x-His or two mutants, FMO1-G17A/G19A-6x-His (a FAD binding mutant) or FMO1-G215A-6x-His (a NADP^+^ binding mutant). 1 mM Pip was co-infiltrated with an *A. tumefaciens* strain carrying *FMO1-6x-His*, *FMO1-G17A/G19A-6x-His*, or *FMO1-G215A-6x-His* and then leaves were harvested 28 hpi. N-OH-Pip and the unknown metabolite were identified as FMO1-dependent signals using an untargeted analysis. Levels represent the mean +/- STD of three biological replicates. Levels reported as zero indicate no detection of metabolites. Pathway represents proposed reactions occurring in *N. benthamiana*. (B) Protein expression of *Arabidopsis* WT and mutant FMO1 enzymes in *N. benthamiana*. Top panel: Immunoblot analysis of *N. benthamiana* total protein extracts containing FMO1-6x-His, FMO1-G17/G19A-6x-His, or FMO1-G215A-6x-His probed with anti-His sera. Bottom panel: Ponceau S-stained Rubisco large subunit (arrow) on membrane is included as a loading control.

### N-OH-Pip treatment of *Arabidopsis* leaves is sufficient to induce SAR

To test if N-OH-Pip treatment alone is sufficient to induce SAR and rescue the SAR deficiency of *fmo1* mutants, we infiltrated three lower leaves of *Arabidopsis* WT and *fmo1* plants with 10 mM MgCl_2_ (Mock) or 10 mM MgCl_2_ containing 1 mM Pip or 1 mM N-OH-Pip. After chemical incubation for 24 hr, one upper leaf of each plant was inoculated with a 1×10^5^ cfu/mL suspension of *Pseudomonas syringae* pathovar *maculicola* strain ES4326 (*Psm* ES4326), a virulent bacterium (Fig. 3A). At three days post inoculation (dpi), bacterial growth was quantified (Fig. 3B), and leaf symptoms were photographed (Fig. 3C). Both WT and *fmo1* plants treated with N-OH-Pip contained significantly less *Psm* ES4326 in infected upper leaves compared to plants treated with Mock or Pip (Fig. 3B). N-OH-Pip treatment also reduced symptom development (*i.e.* leaf yellowing and tissue collapse) (Fig. 3C). These results suggest that treatment of leaves with N-OH-Pip, but not Pip, is sufficient to induce SAR and this this does not require FMO1.

**Figure 3.**
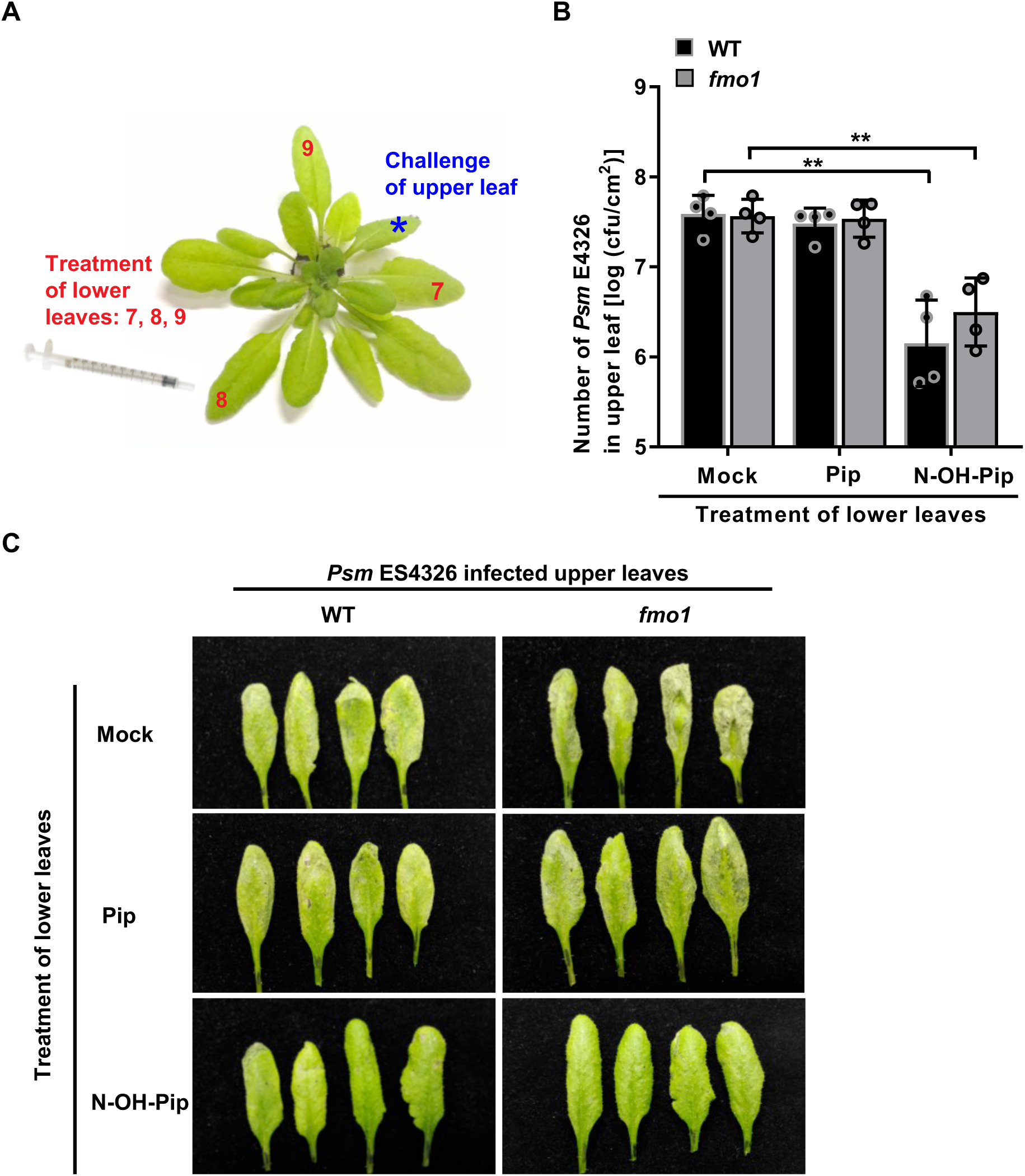
Infiltration of N-OH-Pip into *Arabidopsis* Col-0 WT and *fmo1* leaves inhibits growth of virulent *Psm* ES4326 in distal leaves. (A) Photograph showing a representative plant and leaves used for SAR experiments. Typically, leaf number 7, 8, and 9 were used for chemical treatments (treated lower leaf) and leaf number 11, 12, and 13 were not treated (upper leaf). (B) Titer of *Psm* ES4326 in challenged upper leaves of *Arabidopsis* WT and *fmo1* plants. Three lower leaves of Col-0 WT and *fmo1* were infiltrated with 10 mM MgCl_2_ (Mock) or 10 mM MgCl_2_ containing 1 mM Pip or 1 mM N-OH-Pip. After chemical incubation for 24 hr, one upper leaf of each WT and *fmo1* plants was inoculated with a 1×10^5^ cfu/mL suspension of *Psm* ES4326. The number of *Psm* ES4326 was quantified three days later to measure resistance of leaves. Bars represent the mean +/- STD of four biological replicates. Asterisks denote the significant differences between indicated samples using a one-tailed t-test (** P < 0.01). The experiment was repeated twice with similar results. (C) Symptoms of *Arabidopsis* WT and *fmo1* upper leaves infected with *Psm* ES4326 following chemical treatment described in (B). Leaves were photographed three dpi. The SAR experiment was repeated twice with similar results.

### N-OH-Pip treatment of *Arabidopsis* leaves is sufficient to induce metabolite production and gene transcription associated with SAR

Next we examined the ability of N-OH-Pip to systemically induce the production of SAR-associated metabolites and mRNAs in *Arabidopsis* leaves. Three lower leaves of WT and *fmo1* plants were infiltrated with 10 mM MgCl_2_ (Mock) or 10 mM MgCl_2_ containing 1 mM Pip or 1 mM N-OH-Pip. After 48 hr, the three treated lower leaves and three untreated upper leaves were pooled independently, and then analyzed by LC-MS and qRT-PCR for measurement of metabolites and mRNAs, respectively.

For metabolite profiling, we quantified Pip, N-OH-Pip, N-OGlc-Pip, m/z 100, and two canonical defense metabolites, the phytoalexin camalexin and SA-Glucoside (SA-Glc) (Fig. 4A and 4B). After infiltration of the lower leaves of both WT and *fmo1* plants with N-OH-Pip, we observed accumulation of camalexin and SA-Glc in both lower and upper leaves (Fig. 4B). Our findings are consistent with reports showing that FMO1 is required to stimulate camalexin and SA-Glc synthesis and accumulation in distal, uninfected leaves during SAR [31].

**Figure 4.**
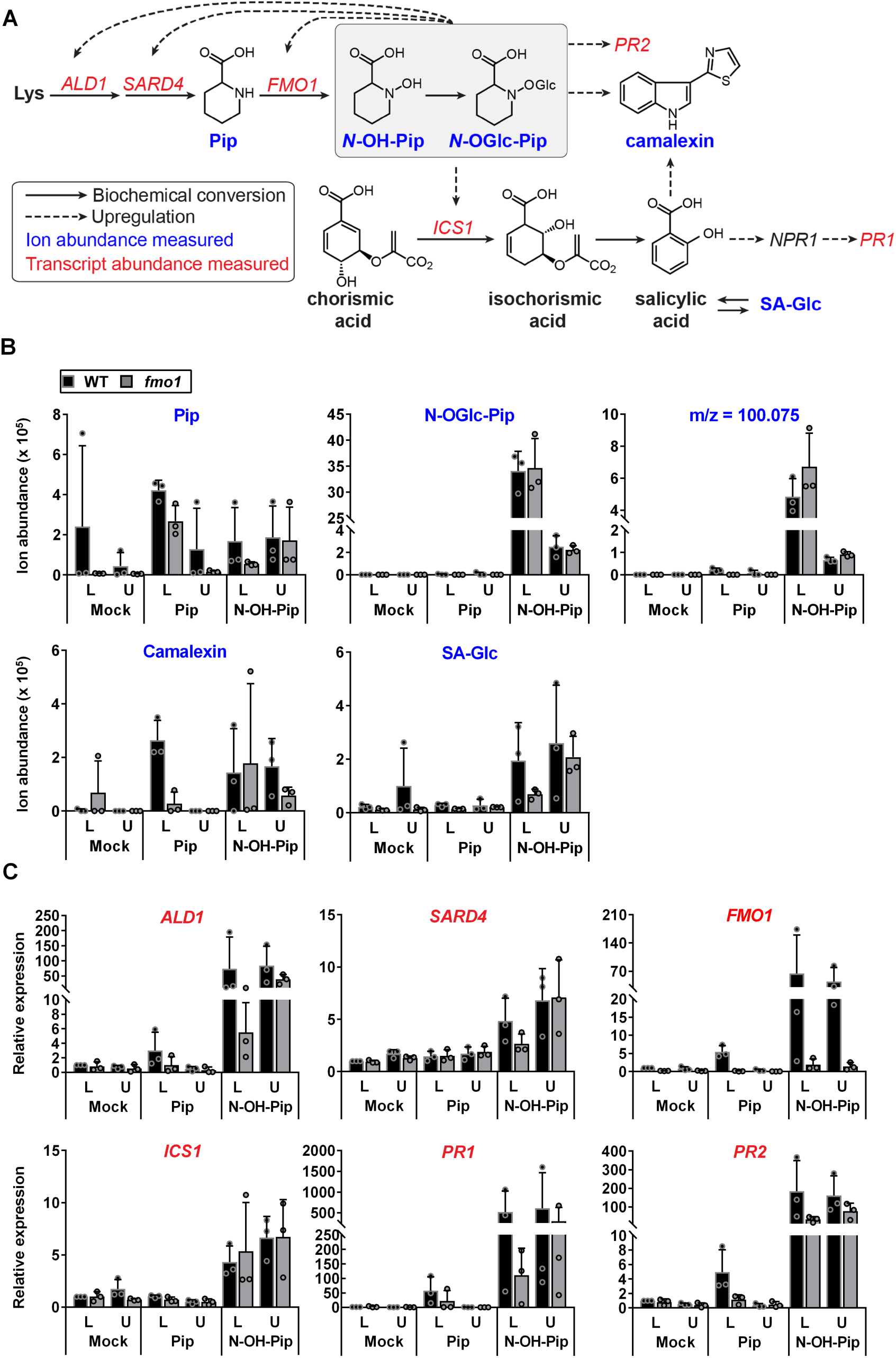
N-OH-Pip induces expression of SAR marker genes and metabolites in treated lower leaves and untreated upper leaves of *Arabidopsis* Col-0 WT and *fmo1* plants. (A) Proposed pathway of metabolite and gene induction in N-OH-Pip-associated SAR. In this study, FMO1 is hypothesized to convert Pip to N-OH-Pip and unknown enzymes convert N-OH-Pip to N-OGlc-Pip, and m/z=100.075. Pip biosynthesis requires ALD1 activity and is partially dependent on SARD4 activity (Navarova et al., 2012; Ding et al., 2016; Hartmann *et* al., 2017). Pip treatment leads to the accumulation of camalexin and *ALD1, SARD4, FMO1, ICS1* and *PR2* mRNAs. ICS1 converts chorismic acid to isochorismic acid, leading to SA production. SA is required for camalexin production in during SAR (Bernsdorff et al., 2016), and SA induces the expression of *NPR1*. NPR1 is a transcription factor that regulates expression of *PR1* and other defense-associated genes. In plants, SA-Glucoside (SA-Glc) is thought to be non-toxic storage form of SA (Dempsey et al., 2011). (B) Ion abundance of Pip, N-OGlc-Pip, m/z=100.075, camalexin and SA-Glc in leaves. Three lower leaves of Col-0 WT and *fmo1* were infiltrated with 10 mM MgCl_2_ (Mock) or 10 mM MgCl_2_ containing 1 mM Pip or 1 mM N-OH-Pip. After chemical incubation for 48 hr, three treated lower (L) leaves and three untreated upper (U) leaves of WT and *fmo1* were analyzed by metabolite profiling and qPCR (C). Three biological replicates were used. The experiment was repeated twice with similar results. (C) Relative expression of SAR marker genes in leaves: *ALD1* (AT2G13810), AGD2-like defense response protein 1; *SARD4* (AT5G52810), SAR deficient 4; *FMO1* (At1g19250), flavin-dependent monooxygenase 1; *ICS1* (At1g74710), isochorismate synthase 1; *PR1* (At2g14610), pathogenesis-related protein 1 and *PR2* (At3g57260), beta-1,3-glucanase 2.

Although we did not detect free N-OH-Pip, we observed accumulation of N-OGlc-Pip and m/z 100 in both lower and upper leaves of all plants (Fig. 4B), suggesting rapid metabolism to these derivatives. These data show that local treatment of purified N-OH-Pip alone, even in the absence of infection, can induce systemic changes in defense metabolite production across the plant and this does not require FMO1. In contrast, the infiltration of lower leaves of WT or *fmo1* plants with Pip did not induce changes in N-OH-Pip metabolites, camalexin, or SA-Glc. Given that *fmo1* plants are unable to produce N-OH-Pip, the detection of N-OH-Pip metabolites (N-OGlc-Pip and m/z 100) in the upper leaves after N-OH-Pip treatment of lower leaves (Fig. 4B) provides evidence that one or more of these metabolites are mobile. Taken together, these data indicate that N-OH-Pip or rapidly formed derivatives move systemically through the plant to induce metabolic changes during SAR.

For transcript profiling, we quantified the mRNA abundance for known SAR marker genes which were reported to be induced in untreated upper leaves of WT plants but not *fmo1* plants in response to bacterial infection (*i.e. Psm* ES4326) of lower leaves [32]. These included genes dependent on SA (*PR1*, pathogenesis-related protein1), partially dependent on SA (*PR2*, pathogenesis-related protein 2; and *PR5*, pathogenesis-related protein 5) or independent of SA (*ALD1*; AGD2-like defense response protein 1; *SAG13*, senescence-associated gene 13; *FMO1*, flavin-dependent monooxygenase 1; and *ICS1*, isochorismate synthase 1). We also monitored *ALD1*, *SARD4* (SAR deficient 4 protein) and *FMO1*, three genes in the proposed N-OH-Pip metabolic pathway (Fig. 1C and 4A).

WT and *fmo1* plants treated with N-OH-Pip accumulated the highest levels of *ICS1* and *PR1* mRNA in treated lower and untreated upper leaves, compared to plants treated with Mock or Pip (Fig. 4C). Similar trends were observed for *ALD1, SARD4, PR2, PR5* and *SAG13* mRNAs (Fig. 4C and Fig. S5). N-OH-Pip triggered accumulation of *ALD1* and *SARD4* mRNA suggests that N-OH-Pip regulates *ALD1* and *SARD4* transcription via a positive feedback loop. Accumulation of Pip in leaves of WT and *fmo1* plants treated with N-OH-Pip implies that N-OH-Pip induction of *ALD1* and *SARD4* transcription leads to increased synthesis of Pip in leaves. In addition, N-OH-Pip treatment increased *FMO1* mRNA levels in WT but not *fmo1* leaves (Fig. 4C). This implies that N-OH-Pip also positively regulates *FMO1* transcription via a positive feedback loop. Notably, induction of SAR gene expression was greater in N-OH-Pip treated plants compared to Pip treated plants (Fig. 4C and Fig. S5), demonstrating that N-OH-Pip is a more bioactive SAR molecule compared to Pip.

Collectively, these data demonstrate that exogenous treatment of N-OH-Pip by leaf infiltration is sufficient to activate metabolite production and SAR-associated gene transcription.

### N-OH-Pip irrigation induces SAR in *Arabidopsis* WT and *fmo1* plants

Previously, Návarová et al. showed that irrigation with Pip was sufficient to recover SAR activity in *ald1* plants, but not in *fmo1* plants [13]. We tested if irrigation with N-OH-Pip could similarly induce SAR in WT and *fmo1* plants. SAR assays were performed as described previously [13] with slight modification. WT and *fmo1* plants were drenched with water, 1 mM Pip, or 1 mM N-OH-Pip by root application. One day later, three lower (local) leaves per plant were inoculated with 10 mM MgCl_2_ (Mock) or a 5 x 10^6^ cfu/mL suspension of *Pst avrRpt2*, an avirulent strain. Two days later, the untreated upper leaves of each plant were challenged with Mock or a 1 x 10^5^ cfu/mL suspension of *Psm* ES4326, a virulent strain. The titer of *Psm* ES4326 in the infected upper leaves was determined at 3 dpi to quantify the impact of Pip and N-OH-Pip treatment on the level of disease resistance. An overview of the SAR experiment is shown in Fig. 5

**Figure 5.**
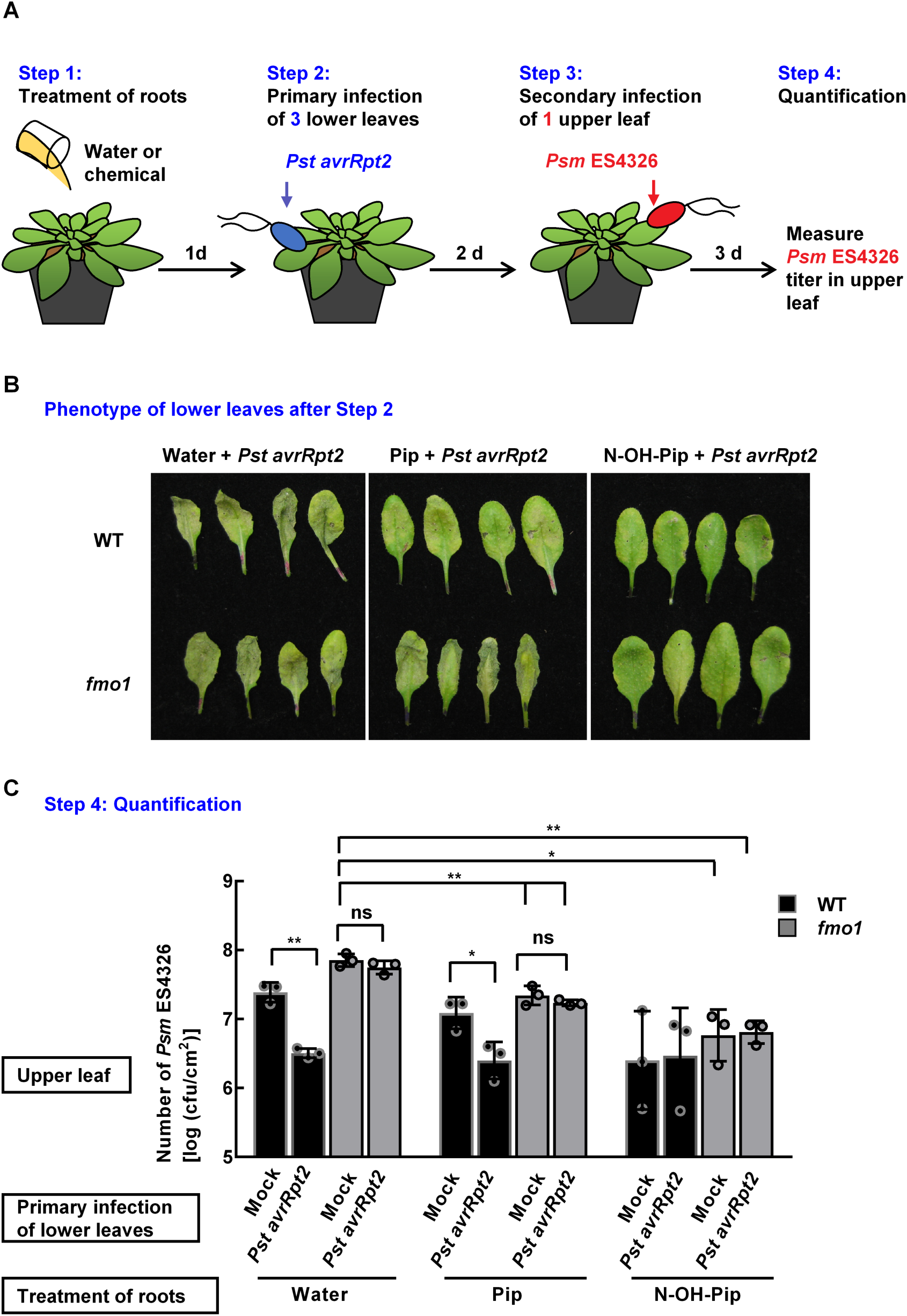
Root application of N-OH-Pip elicits local defense and SAR in *Arabidopsis* Col-0 WT and *fmo1* plants. (A) Experimental design of SAR assay. Each Col-0 WT and *fmo1* plant was treated with 10 mL of water, 1 mM of Pip, or 1 mM of N-OH-Pip by root application. One day later, three lower leaves of each plant were infiltrated with 10 mM MgCl_2_ or a 5×10^6^ cfu/mL suspension of *Pst* DC3000 expressing *avrRpt2* (*Pst avrRpt2*) in 10 mM MgCl_2_. Two days later, one upper leaf of each plant was inoculated with a 1×10^5^ cfu/mL suspension of *Psm* ES4326. The number of *Psm* ES4326 in upper leaves was quantified three days later. (B) Phenotype of lower leaves inoculated with *Pst avrRpt2* after chemical treatment. Phenotypes of four biological replicates were recorded 2 dpi with *Pst avrRpt2*. Similar phenotypes were observed in three independent experiments. (C) Titer of *Psm* ES4326 in upper leaves of Col-0 WT and *fmo1*. Bars represent the mean +/- STD of three biological replicates. Asterisks denote the significant differences between indicated samples using a one-tailed *t-test* (** P < 0.01; * P < 0.05; ns, not significant). The experiment was repeated three times with similar results.

Before challenging upper leaves with *Psm* ES4326, lower leaves infected with *Pst avrRpt2* were yellow (chlorotic) and collapsed by 2 dpi (Fig. 5B). We noticed that these symptoms were significantly delayed in WT and *fmo1* plants irrigated with N-OH-Pip (Fig. 5B). Pip irrigation reduced leaf symptom development in WT plants to a lesser degree, but not in *fmo1* plants (Fig. 5B). Metabolic profiling revealed that N-OGlc-Pip, and m/z 100 were present in *Pst avrRpt2* infected leaves of WT plants irrigated with Pip or N-OH-Pip, and *fmo1* plants irrigated with N-OH-Pip (Fig. S6). These results indicate that N-OH-Pip treatment reduces symptom development of *Arabidopsis* leaves in an *FMO1-*independent manner. The delay of symptom development caused by N-OH-Pip and Pip could be attributed to metabolite-induced defense priming in infected leaves or inherent bactericidal activity of the metabolites.

After challenging upper leaves with *Psm* ES4326, the titer of *Psm* ES4326 in water-treated WT plants infected with *Pst avrRpt2* was lower than that in water-treated WT plants inoculated with Mock (Fig. 5C). These data indicate that *Pst avrRpt2* induces SAR under the conditions tested. By contrast, *Psm* ES4326 titers were similar in water-treated *fmo1* plants inoculated with Mock or *Pst avrRpt2* (Fig. 5C). These data are consistent with previous findings that *fmo1* mutants are deficient in SAR signaling [16, 18].

Irrigating plants with Pip and N-OH-Pip enhanced SAR, albeit to different extents. Pip absorption by roots provided a moderate but significant reduction in *Psm* ES4326 titer in Mock and *Pst avrRpt2* inoculated *fmo1* plants (Fig. 5C). However, the level of protection was less than SAR-induced protection elicited by *Pst avrRpt2* in water-treated WT plants. Interestingly, N-OH-Pip absorption by roots provided the strongest level of protection (Fig. 5C). *Psm* ES4326 titers in Mock and *Pst avrRpt2* inoculated *fmo1* plants were significantly less than those detected in water- or Pip-treated *fmo1* plants. Moreover, N-OH-Pip induced protection in *fmo1* plants was similar to that observed for *Pst avrRpt2* induced protection in WT plants (Fig. 5C). Collectively, these data indicate that N-OH-Pip is a potent SAR-inducing metabolite and irrigation of plants with N-OH-Pip is also sufficient to establish SAR in WT plants and complement the SAR defect of *fmo1* plants.

### N-OH-Pip is not bactericidal

To rule out that N-OH-Pip is toxic to bacterial growth *in planta*, we performed a Minimum Inhibitory Concentration (MIC) assay [33] to measure potential bactericidal activity. Cultures of *Psm* ES4326, *Pst* DC3000 and *Pst avrRpt2* were incubated with different concentrations (0 to 1 mM) of Pip, N-OH-Pip, or SA and then grown at 28°C for 36 h. SA was included as a positive control because it inhibits multiplication of *Pseudomonas aeruginosa* and *Agrobacterium tumefaciens in vitro* [34, 35]. Similar to previous studies, 1 mM SA inhibited the multiplication of *Psm* ES4326, *Pst* DC3000 and *Pst avrRpt2 in vitro* (Fig. S7). However, neither Pip nor N-OH-Pip inhibited *Psm* ES4326, *Pst* DC3000 and *Pst avrRpt2* growth *in vitro*. These data indicate that Pip and N-OH-Pip are not bactericidal.

### N-OH-Pip treatment accelerates the hypersensitive response in *Arabidopsis*

Next we investigated if N-OH-Pip enhances ETI in local, infected leaves by performing electrolyte leakage and HR assays. WT and *fmo1* plants were irrigated with water, 1 mM Pip or 1 mM N-OH-Pip. One day later, six to seven leaves of each plant were inoculated a 3×10^8^ cells/mL suspension of *Pst* DC3000 carrying an empty vector (*Pst* vector) or expressing *avrRpt2* (*Pst avrRpt2*). Ion leakage was quantified at 5 hpi (Fig. 6A) and HR symptoms were recorded at 8 hpi (Fig. 6B).

**Figure 6.**
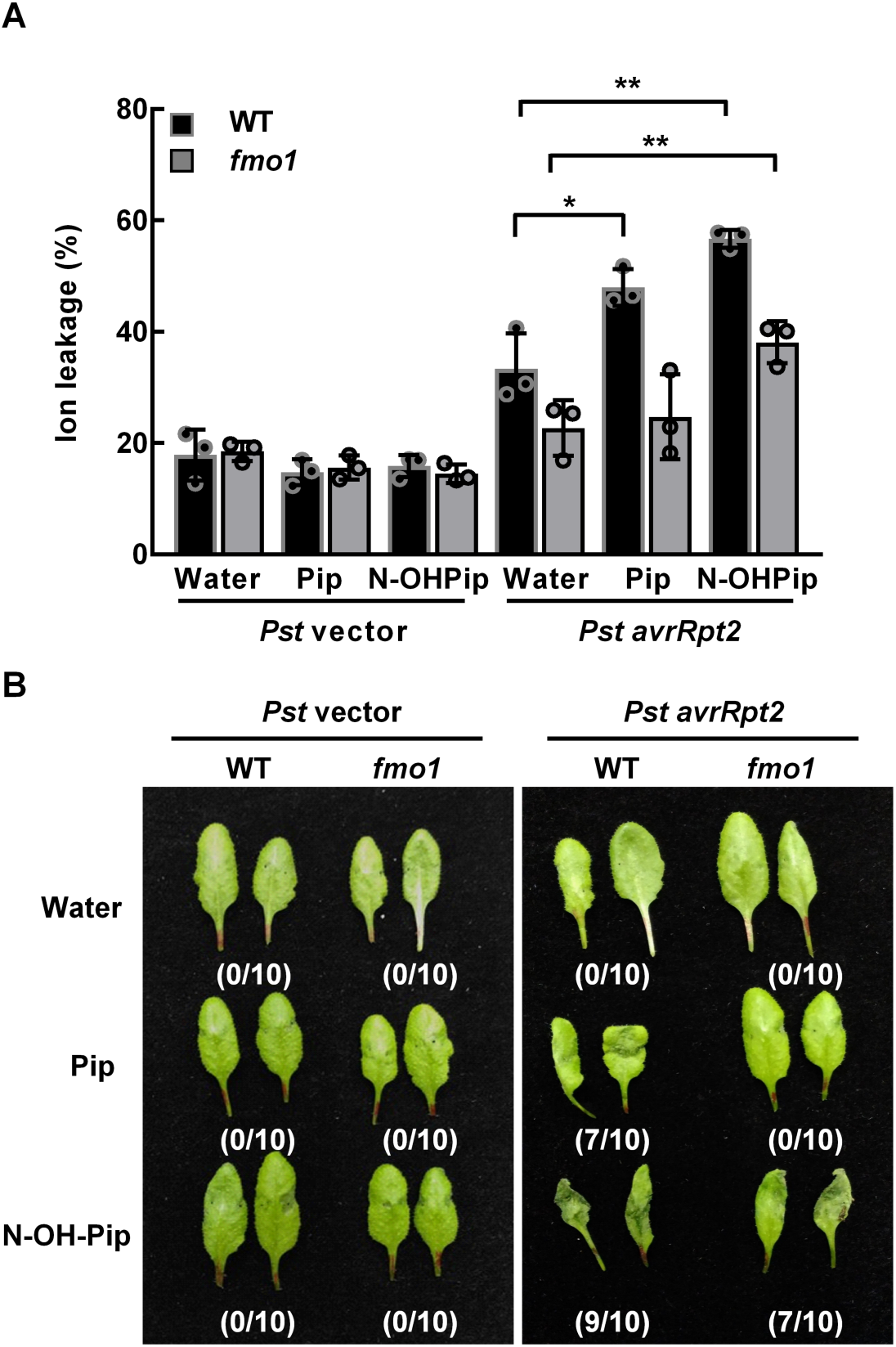
N-OH-Pip accelerates AvrRpt2-elicited HR phenotype in *Arabidopsis* Col-0 WT and *fmo1* plants. Col-0 WT and *fmo1* plant pots were treated with 10 mL of water, 1 mM Pip, or 1 mM N-OH-Pip by root application. Twenty-four hours later, leaves of Col-0 WT and *fmo1* plants were infiltrated with a 3×10^8^ cells/mL suspension of *Pst* DC3000 with an empty vector (*Pst* vector) or *avrRpt2* (*Pst avrRpt2*). (A) Percent ion leakage of inoculated leaves at 5 hours post inoculation (hpi). Bars represent the mean +/- STD of 9 randomly inoculated leaves of three WT or *fmo1* plants. Asterisks denote the significant differences between indicated samples using a one-tailed t-test (** P < 0.01; * P < 0.05). (B) HR phenotypes of inoculated leaves at 8 hpi. Fraction refers to number of leaves showing HR out of 10 randomly inoculated leaves. Experiments were repeated three times with similar results.

WT and *fmo1* leaves infected with *Pst* vector exhibited similar electrolyte leakage (below 20%) when irrigated with water, Pip, and N-OH-Pip (Fig. 6A), and no HR phenotypes were observed for these infected plants at 5 hpi (Fig. 6B). WT plants irrigated with water and infected with *Pst avrRpt2* exhibited more electrolyte leakage at 5 hpi than similarly treated *fmo1* plants (Fig. 6A). HR symptoms were not observed at 8 hpi (Fig. 6B); however, full leaf collapse occurred by 10 hpi. Compared to water irrigation, Pip irrigation resulted in a significant increase in electrolyte leakage from WT leaves infected with *Pst avrRpt2* (Fig. 6A), with most (7/10) of the leaves undergoing HR by 8 hpi (Fig. 6B). No significant changes were detected for similarly treated *fmo1* leaves. Notably, N-OH-Pip irrigation resulted in a significant increase in electrolyte leakage from both WT and *fmo1* leaves infected with *Pst avrRpt2* (Fig. 6A). HR phenotypes for the N-OH-Pip irrigated plants appeared faster and were more severe than those observed for the Pip irrigated plants (Fig. 6B). Taken together, these results show that N-OH-Pip treatment accelerates AvrRpt2-elicited HR in plants when compared with water and Pip treatment, demonstrating that N-OH-Pip treatment also enhances ETI.

## Discussion

FMO1-dependent regulation of defense signaling has emerged as a critical determinant of the establishment and amplification of SAR in *Arabidopsis* [16, 31, 32]. The SAR deficiency of *fmo1* mutants could be due to a failure to produce a priming signal, to recognize defense signal(s), or to transmit defense signal(s) from infected tissue to uninfected tissue. Our discovery that FMO1 produces N-OH-Pip, a bioactive metabolite that is mobile in plants and confers pathogen resistance in systemic tissues, reveals that the key biological function of FMO1 is to produce the priming signal for SAR. The accumulation of N-OH-Pip derived metabolites (N-OGlc-Pip and m/z 100) in untreated, distal leaves of *fmo1* plants following infiltration of N-OH-Pip in local leaves indicates that one or more of these metabolites are transported during SAR.

N-OH-Pip establishment of SAR in *Arabidopsis* is characterized by the induction of transcriptional and metabolic programs in systemic plant tissue. Application of N-OH-Pip to local leaves resulted in the robust activation of several SA-dependent and SA-independent SAR marker genes in distal leaves (Fig. 4C and S5). This is consistent with reports showing that *FMO1* is required for SA-dependent and independent defenses [31, 32]. N-OH-Pip also activated transcription of the N-OH-Pip biosynthetic genes (*ALD1, SARD4* and *FMO1*) in both local and systemic leaves (Fig. 4C), highlighting the importance of transcriptional feedback regulation for amplifying N-OH-Pip-dependent signaling during SAR.

Remarkably, disease symptoms in *Arabidopsis* leaves were dramatically reduced in leaves of bacterially infected plants treated with N-OH-Pip (Fig. 3C and 5B). We speculated that N-OH-Pip might directly affect bacterial growth and/or the ability of bacteria to deliver T3S effectors to plant cells. However, N-OH-Pip was not toxic to pathogen growth in culture (Fig. S7) and did not inhibit the activation of ETI in resistant *Arabidopsis* plants (Fig. 6). These data suggest that N-OH-Pip stimulates or enhances plant-specific processes in infected leaves to prevent their collapse and yellowing. What these physiological processes are remains to be determined.

We also found that N-OH-Pip-treated leaves displayed enhanced resistance in SAR assays (Fig. 5C) and are highly sensitized for ETI (Fig. 6). This suggests that the level of N-OH-Pip in tissues plays a critical role in establishing the magnitude and timing of the disease resistance response. Consistent with our findings, overexpression of FMO1 in *Arabidopsis* resulted in enhanced basal resistance to the oomycete *Hyaloperonospora parasitica* and *Pseudomonas syringae pv. DC3000* (*Pst*), as well as enhanced resistance to *Pst AvrRpt2* [17]. In contrast, loss of FMO1 function resulted in enhanced susceptibility to *Pst* and *H. parasitica* [17, 18].

### Biochemistry of FMO1 and chemistry of Pip hydroxylation

FMOs (flavoprotein monooxygenases) catalyze incorporation of one atom of molecular oxygen onto a nucleophilic substrate [36]. Similar to plant cytochromes P450 that activate and transfer oxygen, FMOs have greatly expanded in plants, suggesting they play critical roles in metabolism and fitness. In *Arabidopsis*, 29 genes encode FMO-like proteins, compared to only 5 in animals [37]. *Arabidopsis* FMOs include the YUCCA group of enzymes, which have been well-studied and are associated with auxin biogenesis, as well as enzymes that S-oxygenate glucosinolates, molecules critical for pathogen defense [37]. FMO1 belongs to a distinct clade of the *Arabidopsis* FMOs, and is notably one of the most overexpressed metabolic genes after pathogen challenge (see Fig. S9). Our discovery that the FMO1 product N-OH-Pip is a bioactive SAR metabolite is further evidence that FMOs have a privileged role in plant defense metabolism. Moreover, it predicts that direct products of FMO1 homologs in other plants may be novel natural products involved in defense priming.

Numerous non-proteinogenic amino acids have been described in biological systems [38], yet N-OH-Pip has not previously been reported. It remains a mystery why N-oxidation of pipecolic acid, a metabolite produced in both plants and bacteria, has evolved as a key step in generating an active signaling molecule for SAR. The hydroxyl amine functionality bears resemblance to the hydroxamic acids required for binding iron in bacterial siderophores, and intriguingly, some bacterial siderophores are now thought to be involved in signaling [39]. Further studies are required to determine if the hydroxylamine substituent of N-OH-Pip has a functional role in its transport, perception, and/or metabolism.

Results from our transient expression experiments in *N. benthamiana* indicate that N-OH-Pip may be unstable *in planta* as FMO1-dependent levels of N-OH-Pip decreased from 28 to 48 hours while levels of m/z 100 increased (Fig S1C). We speculate that O-glycosylation of N-OH-Pip could serve as a stabilization mechanism and that N-OGlc-Pip may the primary storage form of the molecule in *Arabidopsis*. This proposed process is reminiscent of SA glycosylation and storage of SA-Glc in response to pathogen infection [40]. While we cannot definitively rule out that m/z 100 is a product of extraction, its presence could indicate a passive or active attenuation strategy. For example, the level of N-OH-Pip could be controlled via conversion to N-OGlc-Pip for storage or m/z 100 for deactivation. Future work will investigate the precise roles of N-OH-Pip and its derivatives in SAR.

### Presence of N-OH-Pip biosynthesis pathway in other plant species

Because SAR is a broad-spectrum defense strategy shared among many plants, we mined available plant genomes to determine the prevalence of enzymes involved in N-OH-Pip biosynthesis. We tabulated the best BLAST hit between the *Arabidopsis* SARD4, ALD1, and FMO1 proteins and those of sequenced plant genomes, and found that >94% of species analyzed had FMO1 homologs with greater than 50% amino acid identity (Fig. S8). All plants within the Brassicaceae that we evaluated have enzymes with >88% amino acid identity to *Arabidopsis* SARD4, ALD1 and FMO1, and these plants contained the top 5 homologs to the *Arabidopsis* FMO1. In *Arabidopsis*, the closest homolog to FMO1 has only a 74% amino acid identity, and this protein originates from a likely pseudogene of *FMO1* that is missing a large internal coding region. The next best BLAST hit has only a 26% amino acid identity to FMO1. The scarcity of FMO1 homologs in *Arabidopsis* suggests that the homologs in other plants (and especially those of the Brassicaceae) may have the same function. While the N-OH-Pip pathway may be broadly conserved, it is possible that this pathway may have arisen independently in the Brassicaceae and other metabolites may mediate defense signaling outside this plant family.

In conclusion, the discovery of N-OH-Pip as a mobile signal provides critical insight into how plants use small molecules to resist the spread of infection. This work provides new opportunity to address fundamental open questions in SAR biology, including the mechanism of transport, signal perception, and signal attenuation once a heightened defense response is no longer required. The sensitivity of plants to N-OH-Pip treatment also highlights the possibility for translating a chemical or metabolic engineering approach to prime or enhance disease resistance in plants under pathogen pressure.

## Acknowledgements

We thank Russ Li and Russ Stabler for assistance preparing synthetic N-OH-Pip. This work was supported by an HHMI and Simons Foundation Grant 55108565 and NIH DP2 Grant AT008321 (to E.S.S.), DGE-1656518 (to E.C.H.), National Science Foundation IOS-1555957 and Binational Science Foundation Grant 2011069 (to M.B.M.) and Ministry of Science and Technology of Taiwan-105-2917-I-564-093 (to Y.C.C.).

---

**Figure S1.**
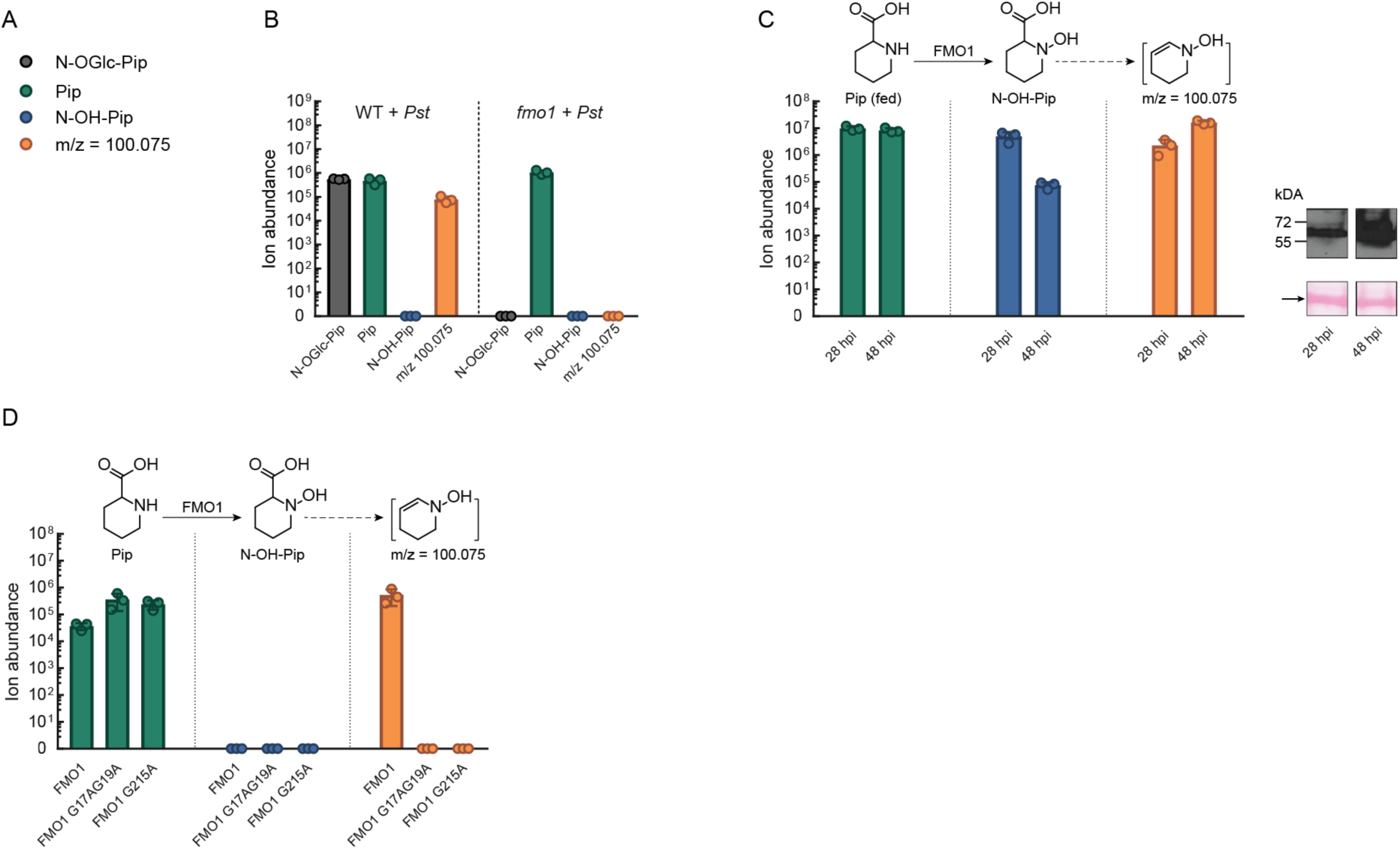
Discovery of N-OH-Pip in *Arabidopsis* and characterization of *Arabidopsis* FMO1 in *Nicotiana benthamiana*. (A) Compounds reported in this figure and their corresponding color key. (B) Ion abundances of N-OGlc-Pip (black), Pip (green), N-OH-Pip (blue), and m/z = 100.075 (orange) in WT and *fmo1 Arabidopsis* seedlings grown hydroponically with a 2×10^8^ cfu/mL suspension of *Pseudomonas syringae* pv. *tomato* DC3000 (*Pst*) as an elicitor. Levels represent the mean +/- STD of six biological replicates. Levels reported as zero indicate no detection of metabolites. (C) Ion abundances of Pip (green), N-OH-Pip (blue), and m/z = 100.075 (orange) in leaves of *N. benthamiana* infiltrated with *Agrobacterium* harboring *FMO1-6x-His* at 28 and 48 hpi. Levels represent the mean +/- STD of three biological replicates. Levels reported as zero indicate no detection of metabolites. Right panel: protein expression of *Arabidopsis* WT FMO1 enzyme in *N. benthamiana* at 28 and 48 hpi. Top: immunoblot analysis of *N. benthamiana* total protein extracts containing FMO1-6x-His probed with anti-His sera. Bottom: ponceau S-stained Rubisco large subunit (arrow) on membrane is included as a loading control. (D) Presence of metabolites during transient expression without supplemental pipecolic acid. Ion abundances of Pip (green), N-OH-Pip (blue), and m/z = 100.075 (orange), in leaves of *N. benthamiana* expressing FMO1-6x-His, FMO1-G17A/G19A-6x-His (a FAD binding site mutant), and FMO1-G215A-6x-His (a NADP^+^ binding site mutant). Levels represent the mean +/- STD of three biological replicates. Levels reported as zero indicate no detection of metabolites.

**Figure S2.**
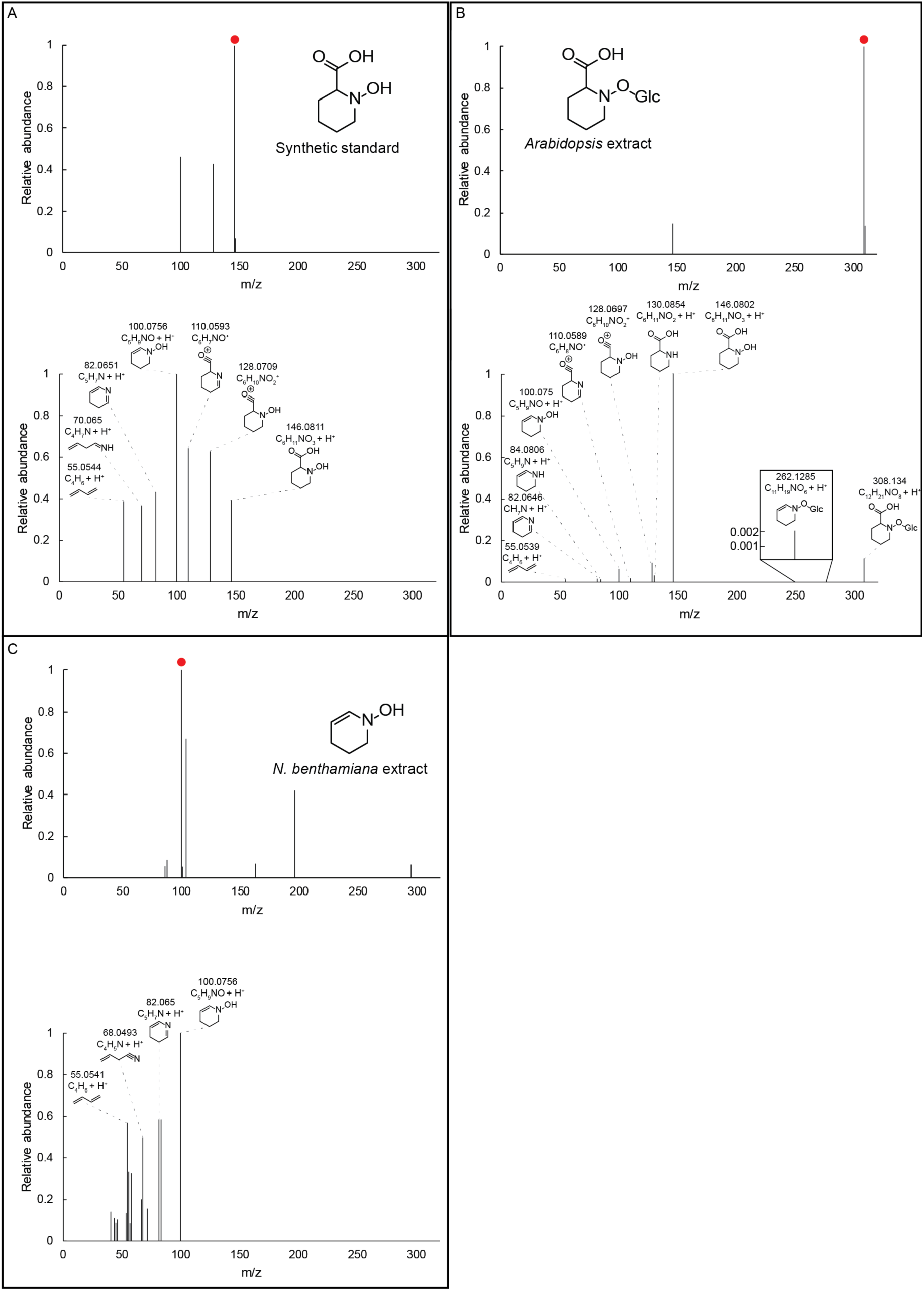
MS/MS spectra for compounds identified in this study. (A) Top: MS of N-OH-Pip synthetic standard at a retention time of 2.2 minutes. Bottom: MS/MS fragmentation pattern of the mass indicated by the red circle. An m/z of 146.081 with a width of 1.3 m/z was used for initial gating. Mass at 146.0811 is the [M+H]^+^ ion of N-OH-Pip. Structures are hypotheses for each fragment ion and masses are the measured mass from the instrument. Only ions with a relative abundance of greater than 5% are shown. (B) Top: MS of N-OGlc-Pip extracted from *Arabidopsis* seedling tissue at a retention time of 2.1 minutes. Bottom: MS/MS fragmentation pattern of the mass indicated by the red circle. An m/z of 308.134 with a width of 1.3 m/z was used for initial gating. Mass at 308.134 is the [M+H]^+^ ion of N-OGlc-Pip. Formulas are hypotheses for each fragment ion and masses are the measured mass from the instrument. With the exception of the ion at m/z = 262, only ions with a relative abundance of greater than 1% are shown. (C) Top: MS of m/z = 100.075 extracted from *N. benthamiana* leaf tissue at a retention time of 1 minute. Bottom: MS/MS fragmentation pattern of the mass indicated by the red circle. An m/z of 100.075 with a width of 1.3 m/z was used for initial gating. Mass at 100.0756 is hypothesized to be the [M+H]^+^ ion of N-hydroxy piperidene. Formulas are hypotheses for each fragment ion and masses are the measured mass from the instrument. Only ions with a relative abundance of greater than 5% are shown.

**Figure S3.**
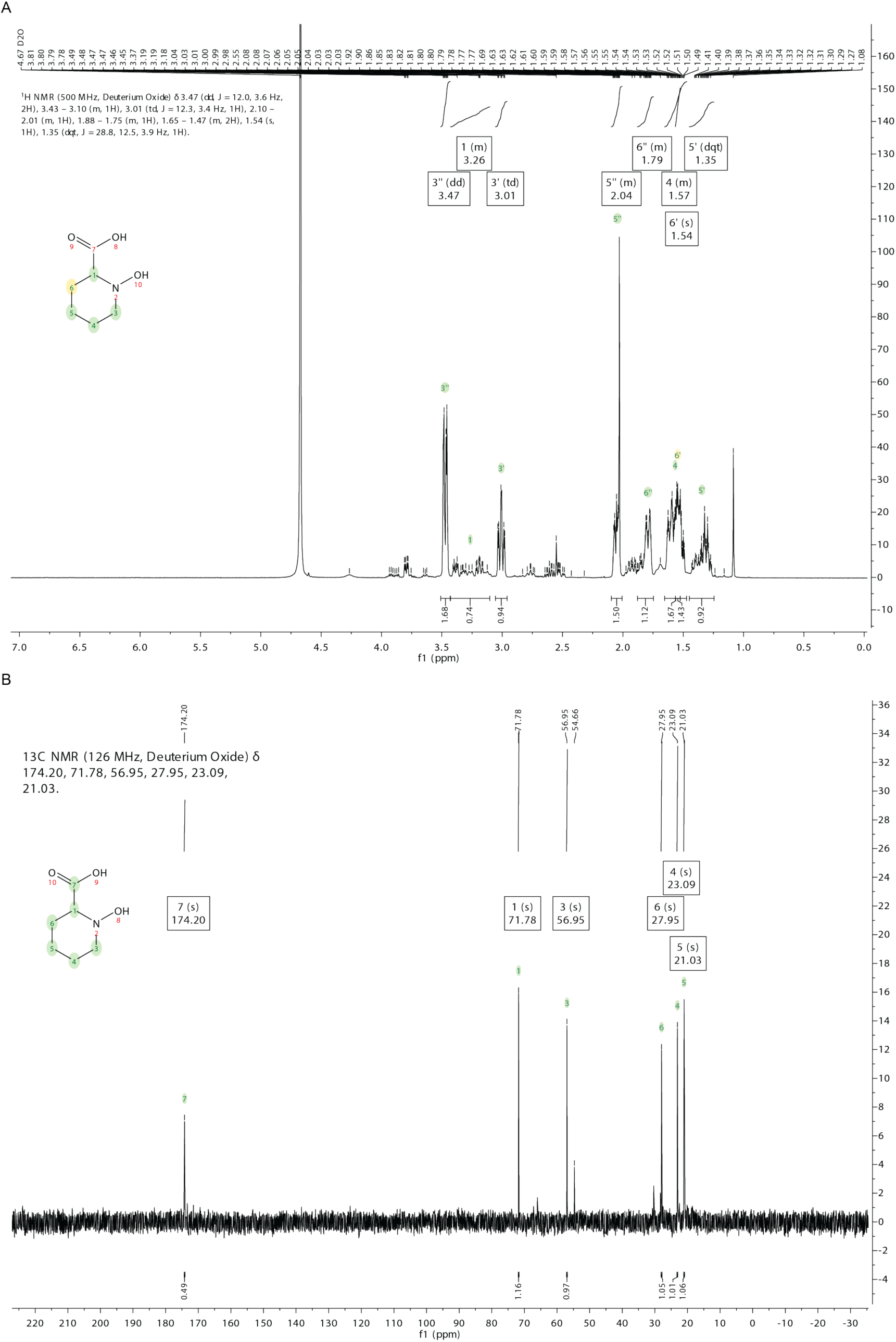
^1^H NMR and ^13^C NMR of synthetic N-OH-Pip standard. (A) Full ^1^H NMR spectrum for synthetic N-OH-Pip standard in D_2_O at ambient temperature and 500 MHz. Chemical shifts and assignment predictions were made using the program MestreNova. (B) ^13^C NMR spectrum of synthetic N-OH-Pip standard in D_2_O at ambient temperature and 126 MHz. Chemical shifts and assignment predictions were made using the program MestreNova.

**Figure S4.**
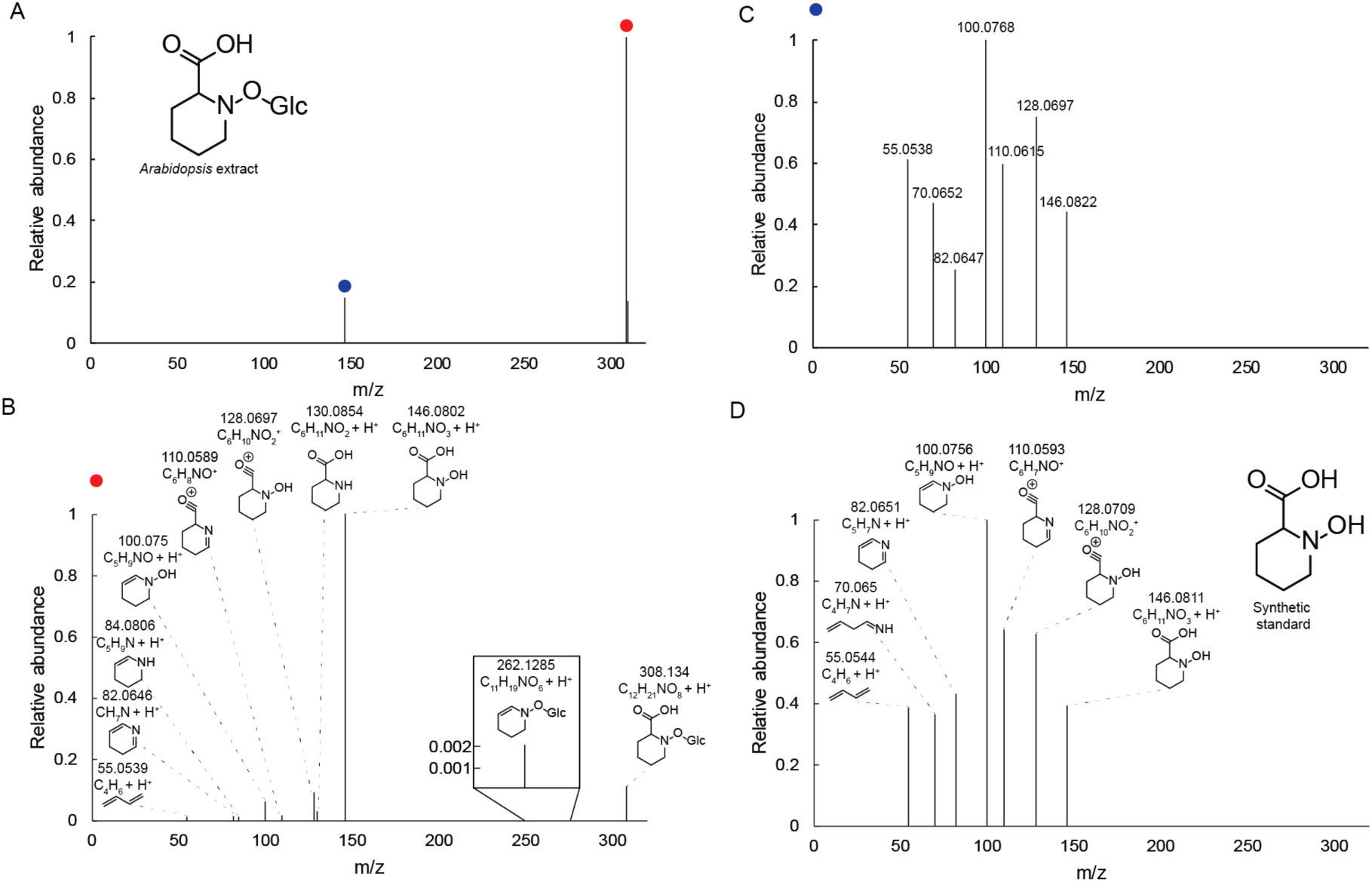
MS/MS fragmentation of N-OGlc-Pip. (A) MS of N-OGlc-Pip extracted from *Arabidopsis* seedling tissue at a retention time of 2.1 minutes. (B) MS/MS fragmentation pattern of the mass indicated by the red circle, hypothesized to be N-OGlc-Pip. An m/z of 308.134 with a width of 1.3 m/z was used for initial gating. Mass at 308.134 is the [M+H]^+^ ion of N-OGlc-Pip. Formulas are hypotheses for each fragment ion and masses are the measured mass from the instrument. Except for the ion at m/z = 262, only ions with a relative abundance of greater than 1% are shown. (C) MS/MS fragmentation pattern of the mass indicated by the blue circle, hypothesized to be an in-source fragment of N-OGlc-Pip to N-OH-Pip. An m/z of 146.081 with a width of 1.3 m/z was used for initial gating. Masses provided are the measured mass from the instrument. Only ions with a relative abundance of greater than 5% are shown. (D) MS/MS fragmentation pattern of N-OH-Pip synthetic standard. An m/z of 146.081 with a width of 1.3 m/z was used for initial gating. Structures are hypotheses for each fragment ion and masses are the measured mass from the instrument. Only ions with a relative abundance of greater than 5% are shown.

**Figure S5.**
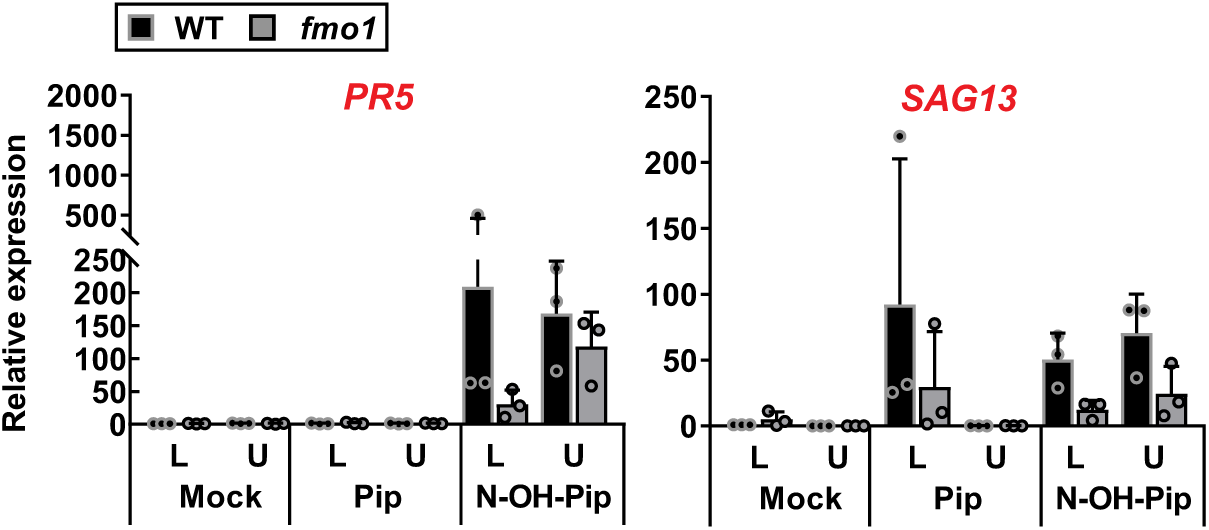
N-OH-Pip induces expression of SAR marker genes in treated lower leaves and untreated upper leaves of *Arabidopsis* Col-0 WT and *fmo1* plants (related to Figure 4C) Three lower leaves of WT and *fmo1* were infiltrated with 10 mM MgCl_2_ (Mock) or 10 mM MgCl_2_ containing 1 mM Pip or 1 mM N-OH-Pip. Forty-eight hr later, three treated lower (L) leaves and three untreated upper (U) leaves analyzed by qPCR. Relative expression of SAR marker genes in leaves. *PR5* (AT1G75040), pathogenesis-related protein 5; *SAG13* (AT2G29350) senescence-associated gene 13), a short-chain alcohol dehydrogenase. Three biological replicates were used in this experiment. The experiment was repeated twice with similar results.

**Figure S6.**
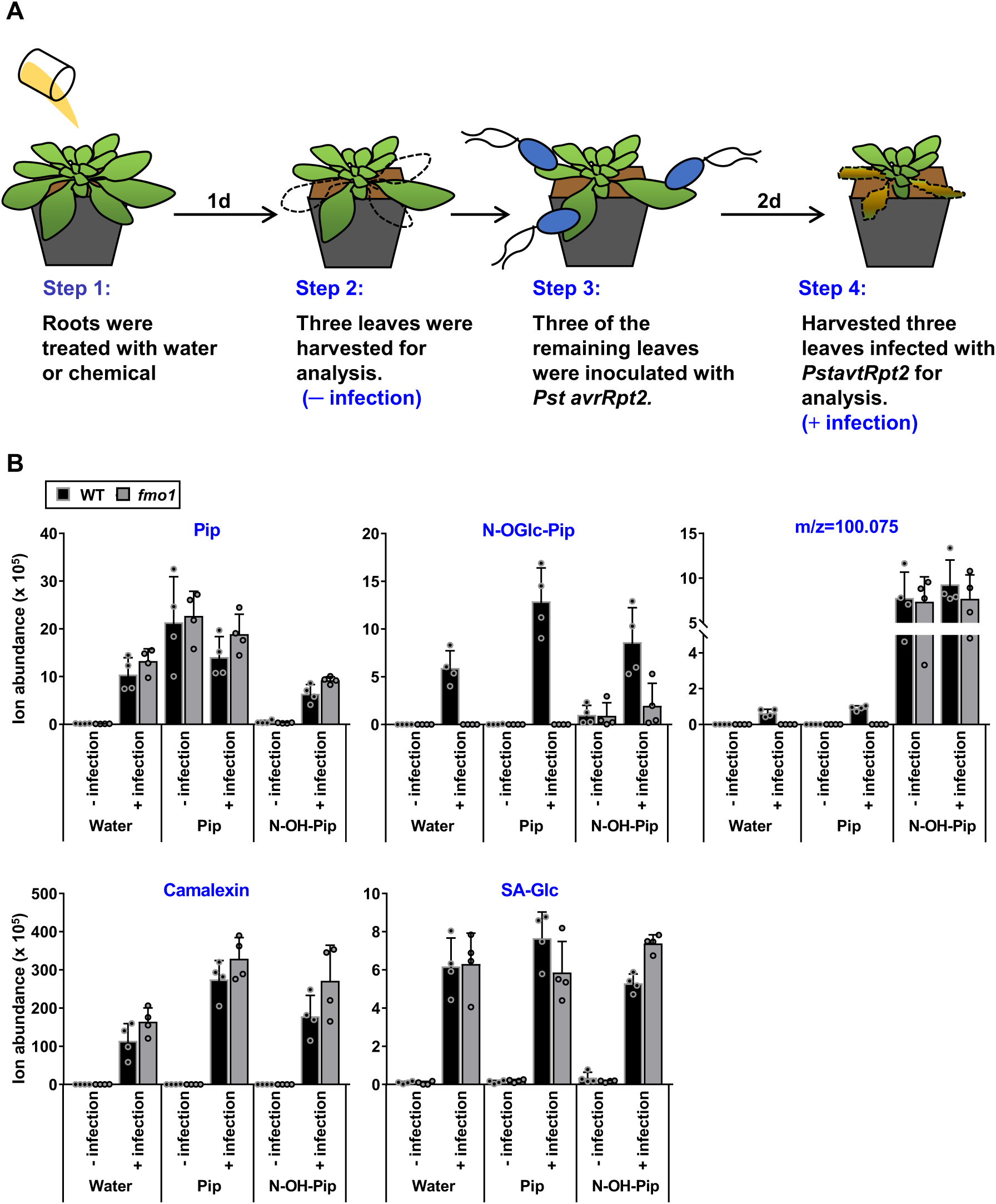
Metabolite profiling of leaves of chemically treated *Arabidopsis* WT and *fmo1* plants before and after *Pst avrRpt2* infection (related to Figure 5B) Each Col-0 WT and *fmo1* plant was treated with 10 mL of water, 1 mM of Pip, or 1 mM of N-OH-Pip by root application. After chemical incubation for 24 hr, three random leaves of each plant were harvested (- infection). Three of the remaining leaves of each plant were infiltrated with a 5×10^6^ cfu/mL suspension of *Pst avrRpt2* in 10 mM MgCl_2_ and harvested at 48 hpi (+ infection). Ion abundance of Pip, N-OGlc-Pip, m/z = 100.075, camalexin and SA-Glc in leaves. Four biological replicates were used in this experiment. The experiment was repeated twice with similar results.

**Figure S7.**
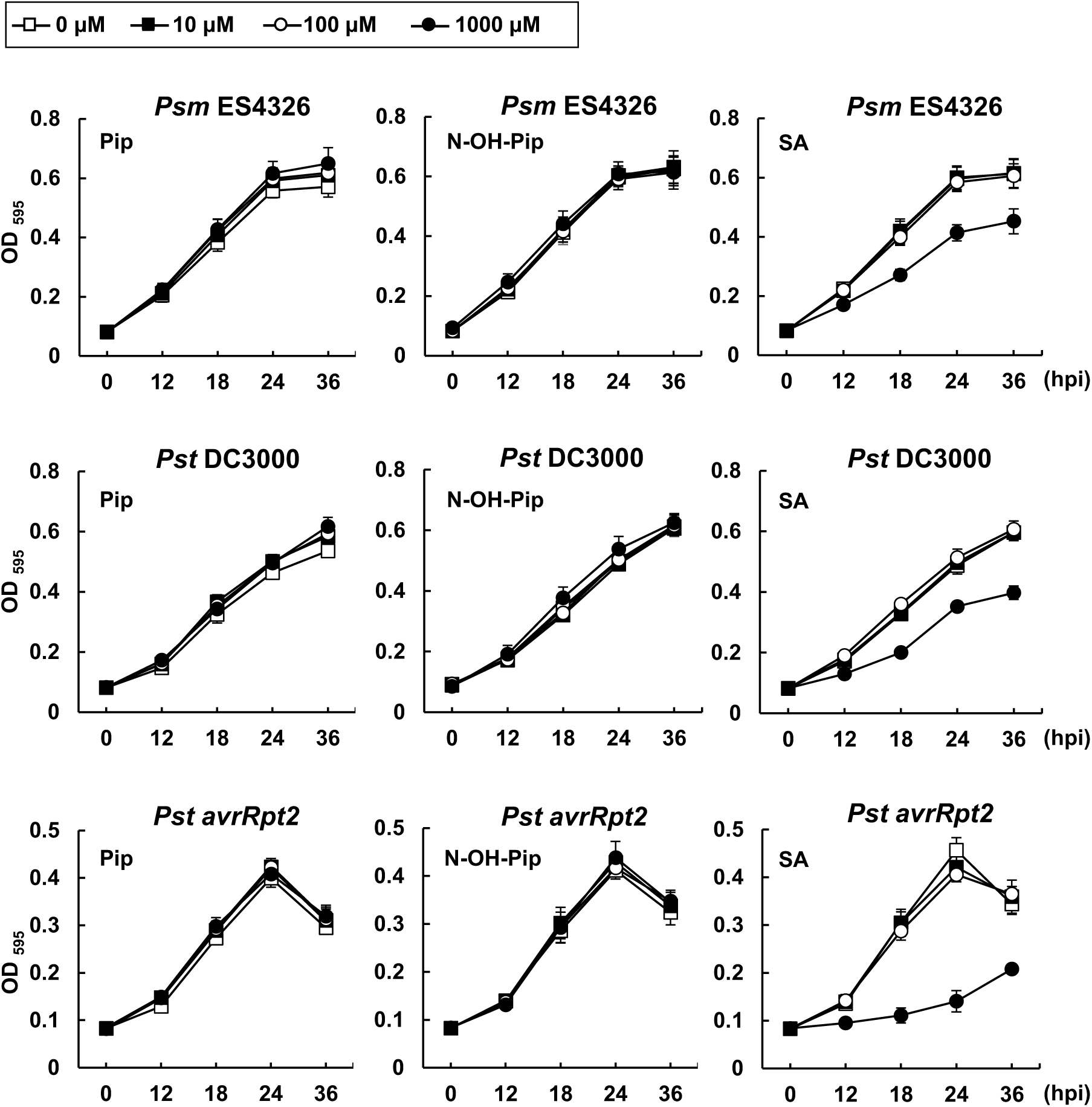
Impact of Pip, N-OH-Pip and salicylic acid (SA) on *Psm* ES4326*, Pst* DC3000 and *Pst avrRpt2* growth. Different concentrations of Pip, N-OH-Pip or SA were mixed with a 10^6^ cfu/mL suspension of *Psm* ES4326, *Pst* DC3000 or *Pst avrRpt2* in 96-well microplates and incubated at 28°C. OD595 nm measurements of the *Psm* ES4326*, Pst* DC3000 and *Pst avrRpt2* cultures were recorded at 0, 12, 24 and 36 hr later. Bars represent the mean +/- STD of four biological replicates. Similar results were observed in three independent experiments for *Psm* ES4326 and *Pst* DC3000 and in two independent experiments for *Pst avrRpt2*.

**Figure S8.**
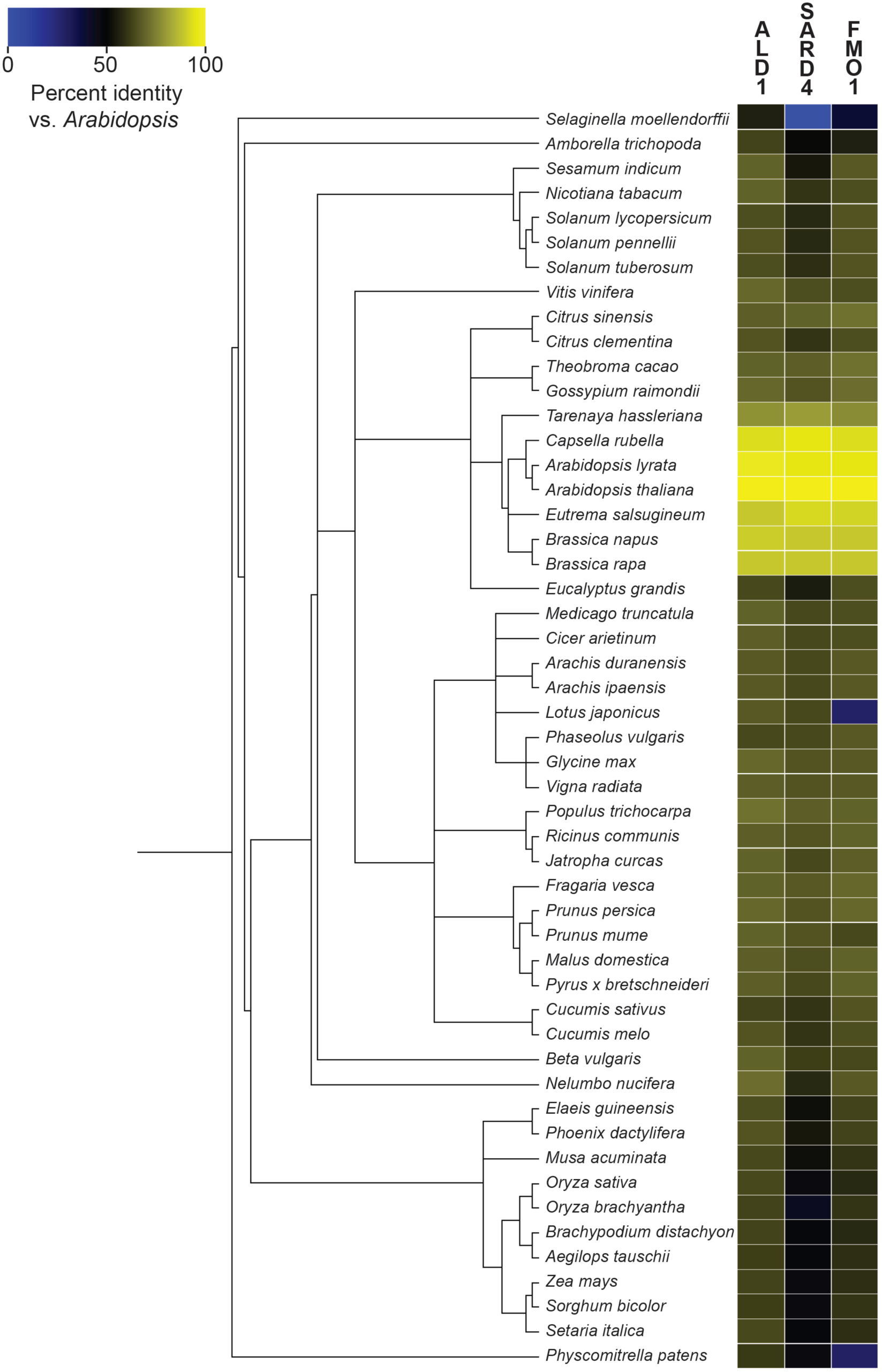
Homologs of N-OH-Pip biosynthetic proteins in genome-sequenced plants. Phylogenetic tree of sequenced plant genomes created using the PhyloT tree generator (http://phylot.biobyte.de/index.html). Heatmap shows the percent amino acid identity of the best BLAST (blastp) hit between the *Arabidopsis* ALD1, SARD4, and FMO1 proteins and those of the respective plant proteome.

**Figure S9.**
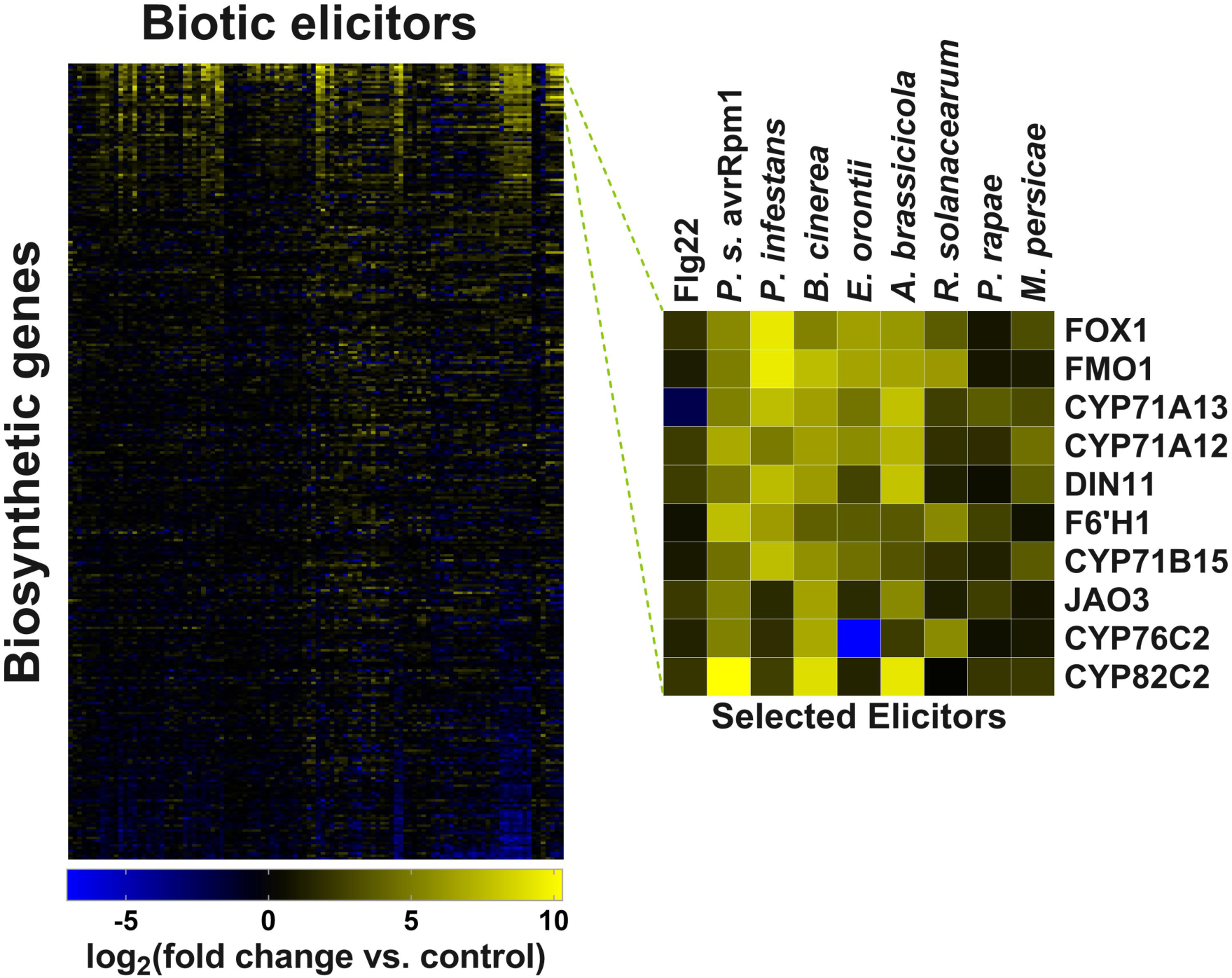
Gene expression analysis of *Arabidopsis* CYP450s, 2-ODDs, and Flavin-dependent oxidative enzymes. Relative gene expression levels of *Arabidopsis* CYP450s, 2-ODDs, and Flavin-dependent oxidative enzymes under various biotic elicitors. The enlarged heatmap shows the top 10 overexpressed genes in response to biotic elicitors after sorting by relative expression over all conditions.

**Table S1.** Strains Used in This Study.

| **Strains** | **Growth conditions** |
| --- | --- |
| *Pseudomonas syringae* pv. *maculicola* strain ES4326<br>**(Psm ES4326)** | LB, NYGB or NYGA containing 100 µg/mL rifampin; 28°C |
| *Pseudomonas syringae* pv. *tomato* DC3000<br>**(Pst DC3000)** | LB or LA containing 100 µg/mL rifampin; 30°C |
| *Pseudomonas syringae* pv. *tomato* carrying pVSP61 + *avrRpt2*<br>**(Pst avrRpt2)** | NYGB or NYGA containing 100 µg/mL rifampin and 50 µg/mL kanamycin; 28°C |
| *Pseudomonas syringae* pv. *tomato* carrying pVSP61 vector<br>**(Pst vector)** | NYGB or NYGA containing 100 µg/mL rifampin and 50 µg/mL kanamycin; 28°C |
| *Escherichia coli* strain DH5 alpha<br>**(E. coli DH5 alpha)** | LB or LA; 37°C |
| *Agrobacterium tumefaciens* strain C58C1<br>**(A. tumefaciens C58C1, pCH32)** | LB or LA containing 100 µg/mL rifampin and 5 µg/mL tetracycline; 28°C |

Lysogeny broth medium (LB; pH 7.0, 1% tryptone, 0.5% yeast extract, 171 mM NaCl); LA (LB supplemented with 1.5% w/v agar); Nutrient yeast glycerol medium (NYGB; pH 7.0, 0.5% peptone, 0.3% yeast extract, 2% glycerol); NYGA (NYGB supplemented with 1.5% w/v agar).

**Table S2.**
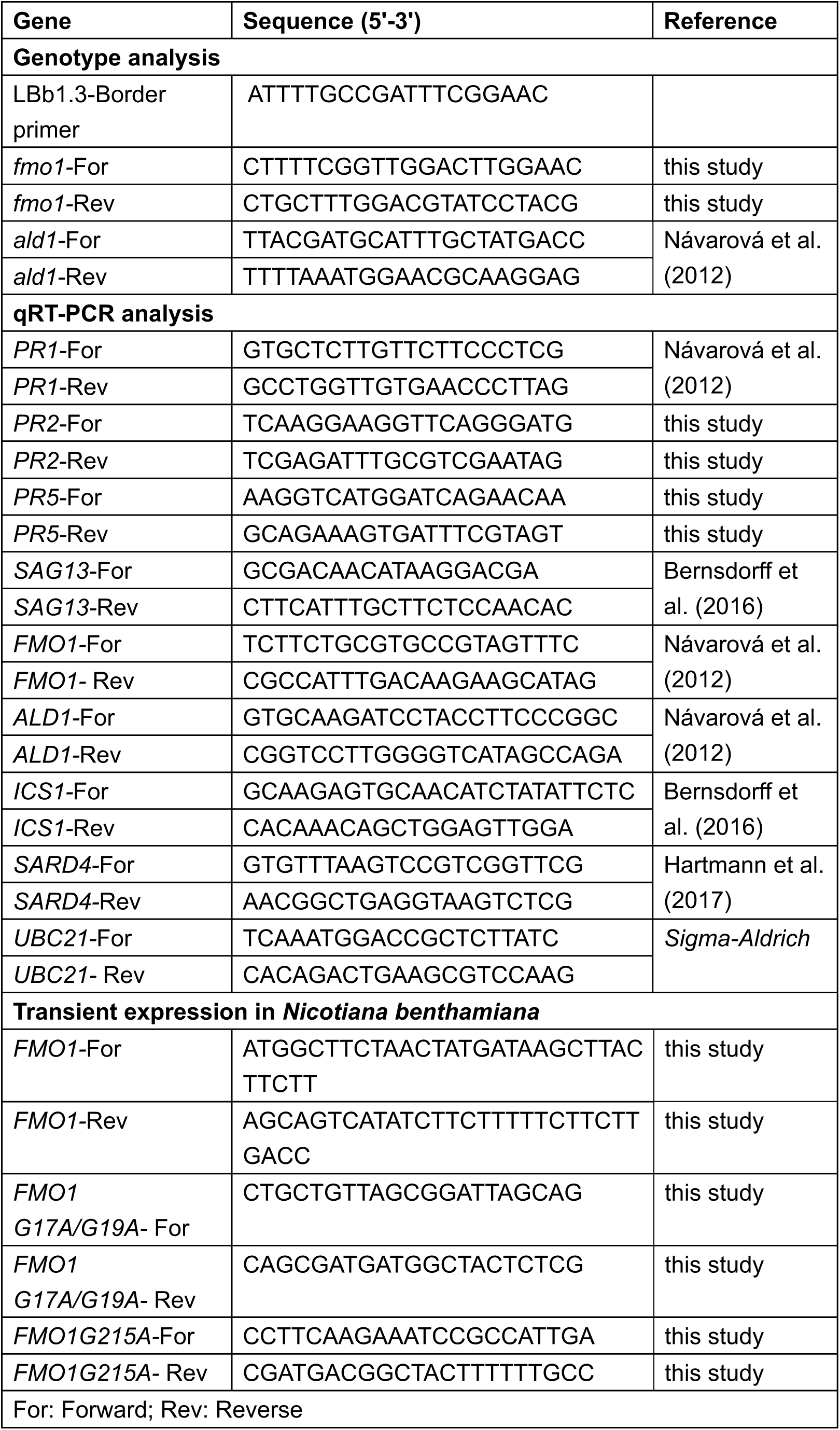
Primers Used in This Study.

## Methods

### Transcriptomics

*Arabidopsis* microarray datasets were downloaded from the NASCArrays database (Craigon et al., 2004). Data from experiments 41, 56, 59, 120, 122, 123, 167, 168, 169, 321, 330, 398, 403, 414, 415, 447, 454, 455, 463, and 468 were used for analysis (indexed experiments can be found at http://arabidopsis.info/affy/link_to_iplant.html). Log-scaled gene expression ratios were calculated as previously (Rajniak et al., 2015) using the following formula:

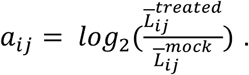

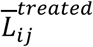 is the mean transcript abundance for gene probe *i* under treatment condition *j* 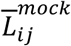 and is the corresponding mock treatment that was used for normalization across experimental conditions.

### Bacterial Strains and Growth Conditions

*Pseudomonas syringae* pv*. maculicola* strain ES4326 (*Psm* ES4326), *Pseudomonas syringae* pv*. tomato* DC3000 (*Pst* DC3000), *Pseudomonas syringae* pv*. tomato* carrying pVSP61 + *avrRpt2* (*Pst avrRpt2*), *Pseudomonas syringae* pv*. tomato* carrying pVSP61 vector (*Pst* vector), *Escherichia coli* strain DH5 alpha, and *Agrobacterium tumefaciens* strain C58C1 pCH32 were used in this study. Growth conditions for all bacterial strains are described in Table S1.

### Plant Materials and Growth Conditions

*Arabidopsis thaliana* ecotype Col-0 (wild-type, WT) and *fmo1* (SALK_026163) plants were grown in a controlled growth chamber (22°C, 80% RH, 103 µmol/m²/s of light intensity) on a 10-hr light /14-hr dark cycle. Genotypes were confirmed by PCR using *FMO1* and *ALD1* gene-specific primers and a T-DNA-specific primer (Table S2). Four to five-week-old adult plants were used for SAR assays, qRT-PCR and metabolite profiling. For seedling hydroponics experiments, *Arabidopsis* seeds were sterilized by suspending seeds in 50% ethanol for 1 min, washing three times in sterile water, suspending in 50% bleach for 10 min, and washing three more times in sterile water. Seeds were resuspended in 1x Murashige-Skoog (MS) medium with vitamins (PhytoTechnology Laboratories) (pH 5.7) and vernalized for 48 hr at 4°C. 15 +/- 1 seeds were placed into 3 mL MS medium + 5 g/L sucrose in wells of a 6-well microtiter plate. Plates were sealed with Micropore tape (3M) and grown in a chamber at 50% humidity, 22°C, and 100 µmol/m^2^/s photon flux under a 16-hr light / 8-hr dark cycle. After 1 week of growth, spent medium was replaced with 3 mL of fresh MS medium + 5 g/L sucrose and plants were grown for 1 additional week. *Nicotiana benthamiana* plants were grown in soil on a growth shelf with a 16 hr light cycle for 5 weeks.

### Elicitation Methods for Seedling Hydroponics Experiments

For bacterial elicitation, *Pst* was grown on LB agar plates at 30°C. A single colony was picked and grown to exponential phase in liquid LB at 30°C. Cells were then centrifuged at >13,000 x *g* and resuspended in sterile 1x MS media to an OD_600_ of 0.2. 70 µl of the *Pst* suspension was added to each seedling-containing well for elicitation. Plants were elicited 24 h prior to sample harvest.

### Plant Extraction for Metabolite Profiling

*Arabidopsis* seedlings were lyophilized to dryness and ground using a ball mill (Retsch MM 400). A single 5 mm steel ball was added to each sample and samples were shaken at 25 Hz for 2 min. Samples were resuspended in 10 mL 80:20 MeOH:H_2_O per gram wet tissue and heated at 65°C for 10 min. Samples were filtered through 0.45 µm PTFE filters and analyzed via liquid chromatography mass spectrometry (LC-MS). Adult leaf tissue was flash frozen and ground in liquid nitrogen using a mortar and pestle. Samples were resuspended in 10 mL 80:20 MeOH:H_2_O per gram wet tissue and heated at 65°C for 10 min. Extracts were filtered through 0.45 µm PTFE filters and analyzed via LC-MS.

### LC-MS Methods

N-OGlc-Pip, SA-Glc, and camalexin were measured using reverse-phase chromatography on an Agilent 1260 HPLC coupled to an Agilent 6520 Q-TOF ESI mass spectrometer as previously (Rajniak et al., 2015). A 5 μm, 2 × 100 mm Gemini NX-C18 column (Phenomenex) was used for separation. The mobile phases were A (water with 0.1% formic acid (FA)) and B (acetonitrile (ACN) with 0.1% FA) and the following gradient was implemented at a flow rate of 0.4 mL/min (percentages indicate percent B): 0-30 min (3-50%), 30-45 min (50-97%), 45-50 min (97%), 50-51 min (97-3%), 51-55 min (3%). The MS was run in positive ion mode with conditions as used previously (Rajniak et al., 2015). For MS/MS analysis, a fragmentor voltage of 10 V was used with an m/z window of 1.3.

Pip, N-OH-Pip, and hypothesized N-hydroxy piperidene were measured using hydrophilic interaction chromatography on an Agilent 1290 UHPLC coupled to an Agilent 6545 Q-TOF ESI mass spectrometer. A 130Å, 1.7 µm, 2.1 mm X 50 mm Acquity UPLC BEH Amide column (Waters) was used for separation. The mobile phases were A (water, 10 mM ammonium formate, 0.125% FA) and B (95% ACN, 10 mM ammonium formate, 0.125% FA), and the following gradient was implemented at a flow rate of 0.6 mL/min (percentages indicate percent B): 0-1.5 min (100%), 1.5-6 min (100-40%), 6-8 min (40%), 8-8.5 min (40-30%), 8.5-9.5 min (30-100%), 9.5-12 min (100%). The MS was run in positive ion mode with the following parameters: mass range: 30-1700 m/z; drying gas: 250°C, 12 L/min; nebulizer: 10 psig; capillary: 3500V; fragmentor: 140 V; skimmer: 65 V; octupole 1 RF Vpp: 750 V; 333.3 ms/spectrum. For MS/MS analysis, a fragmentor voltage of 10 V was used with an m/z window of 1.3 for N-OH-Pip and a fragmentor voltage of 20 V with an m/z window of 1.3 was used for hypothesized N-hydroxy piperidene. The initial 0.5 min of each run was sent to waste to avoid salt contamination of the MS.

### LC-MS Data Analysis

Targeted analysis of metabolites was performed using MassHunter Qualitative Analysis software (Agilent). To obtain ion abundances, ion chromatograms of the major product ion were extracted with a 20 ppm error. Integration was then performed on the appropriate peaks using the built-in peak integration tool in MassHunter. Ions measured in this study can be found in Table E1.

Untargeted metabolite analysis was performed using the R-packages XCMS (Smith et al., 2006) and CAMERA (Kuhl et al., 2012). To input files into XCMS, files were first converted from the Agilent format (.d) to .mzXML files using MSConvert (Kessner et al., 2008). Samples were grouped by experimental condition and analyzed using a standard XMCS and CAMERA workflow shown below:

~~~
##########################################
##Data analysis using xcms
##########################################

#Load xcms and set working directory

library(xcms)
library(CAMERA)
setwd(“YOUR WORKING DIRECTORY”)
options(max.print = 50)

#####################################
##Data analysis
#####################################

pk <- xcmsSet() #Make peak list for every file
peaklisted <- xcmsSet(fwhm = 5, max=30) #If setting parameters for fwhm and max
pk_align <- group(pk) #Align peaks across multiple files
pk_cal <- retcor(pk_align, family=“s”, plottype=“m”) #Recalibrates time axis for all files
pk_cal_g <- group(pk_cal, bw=10) #Regroup calibrated peaks with higher stringency
pk_final <- fillPeaks.chrom(pk_cal_g) #Fills in no peaks with baseline to help statistical analysis

#Generate final report. Must change output names to match folders
Reporttab <- diffreport(pk_final, “FMO1”, “GFP”, “sig_peaks”, 100)

#Do CAMERA analysis

xsa <- xsAnnotate(pk_final)
#Group after RT value of the xcms grouped peak
xsaF <- groupFWHM(xsa, perfwhm=0.6)
#Verify grouping
xsaC <- groupCorr(xsaF)
#Annotate isotopes, could be done before groupCorr
xsaFI <- findIsotopes(xsaC)
#Annotate adducts
xsaFA <- findAdducts(xsaFI, polarity=“positive”)
#Get final peaktable and store on harddrive
write.csv(getPeaklist(xsaFA),file=“result_CAMERA.csv”)
~~~

The outputs from the XCMS workflow were filtered, and only peaks with an average intensity of >50,000, a *p*-value between experimental groups of <0.05, and a fold change between experimental groups of >5 were considered. Peaks were then further filtered by visualization and irregular peaks were thrown out. The remaining peaks were grouped by retention time using CAMERA, which identifies isotopes and adducts that originate from the same molecule.

### N-OH-Pip Chemical Synthesis

N-OH-Pip was synthesized from L-pipecolic acid (98% purity, Oakwood Chemical) using a modified protocol (Nagasawa et al., 1972). To begin, 10.1 g L-pipecolic acid was added to a cooled solution of 4.93 g 88% pure KOH (1 equivalent). 5.58 mL acrylonitrile (1.1 equivalents) was then added drop wise to the solution over 5 min. The solution was stirred for 1.5 h in an ice bath and another 1.5 h at room temperature. Then, the pH of the solution was adjusted to 6.6 with 12 N HCl to quench the KOH. A rotary evaporator was used to evaporate the solvent. 400 mL acetone was added to the residue, and the solution was brought to a boil. After several min of boiling, the solution was filtered and a rotary evaporator was used to remove approximately half of the solvent. Then, the residue from the filtration and fresh acetone was added back into the remaining filtrate, and the solution was brought to a boil, filtered and set on the rotary evaporator. The cycle of heating the solution and filtering was repeated five times. 200 mL filtrate from the 5th cycle was then stored at -20°C overnight to recrystallize.

3.78 g crystallized product (2-cyanoethyl-pipecolic acid) was mixed with 60 mL MeOH and 4.80 g 70% meta-chloroperoxybenzoic acid (mCPBA) (1 equivalent) was added to 20 mL MeOH. The mCPBA solution was added drop wise to the cooled 2-cyanoethyl- pipecolic acid slurry over 30 min. The slurry was stirred for an additional h in an ice bath, and as the reaction progressed, the slurry dissolved into solution. Then, 300 mL precooled diethyl ether was added to the reaction and the mixture was stored at -20°C overnight to recrystallize.

0.5 g crystallized product (2-cyanoethyl-pipecolic acid oxide) was dissolved in 150 mL acetone in a 250 mL flask with a short path distillation head. Acetone was slowly distilled drop by drop for 3 h, and fresh acetone was periodically added to keep the original volume. Then, a rotary evaporator was used to evaporate the majority of the solvent and the remainder was evaporated to dryness under reduced pressure.

^1^H and ^13^C NMR spectra were taken of a 25 mM solution of N-OH-Pip in D_2_O with a Varian Inova 500 NMR spectrometer. The following parameters were used for ^1^H NMR spectra: temperature: ambient; probe: 5mm PFG switchable; scan number: 16; receiver gain: 40; relaxation delay: 0; pulse width: 8; frequency: 499.75 Hz. The following parameters were used for ^13^C NMR spectra: temperature: ambient; probe: 5mm PFG switchable; scan number: 120; receiver gain: 54; relaxation delay: 0.5; pulse width: 7; frequency: 125.67 Hz.

### Chemical Treatment of Leaves for Bacterial Growth Assay

The process of leaf numbering was performed according to (Farmer et al., 2013; Mousavi et al., 2013). Three lower leaves (leaf number 7-9) of WT and *fmo1 Arabidopsis* plants (4- to 5-week old) were infiltrated with 10 mM MgCl_2_, 1 mM Pip in 10 mM MgCl_2_, or 1 mM N-OH-Pip in 10 mM MgCl_2_. After 24 hr, one untreated upper leaf (leaf number 11, 12 or 13) of each plant was inoculated with a 1×10^5^ cfu/mL suspension of *Psm* ES4326. The inoculated plants were then covered with a dome to maintain humidity. The titer of *Psm* ES4326 in the upper leaves was quantified at 3 days post-infiltration (dpi) by homogenizing leaves discs in 1 mL 10 mM MgCl_2_, plating appropriate dilutions on NYGA medium with rifampicin (100 µg/mL), incubating plates at 28°C for 2 days, and then counting bacterial colonies. Four biological repeats were performed per treatment in two independent experiments.

### Chemical Treatment of Leaves for qRT-PCR and Metabolite Profiling

Three lower leaves (leaf number 7-9) of WT and *fmo1 Arabidopsis* plants (4- to 5-week old) were infiltrated with 10 mM MgCl_2_, 1 mM Pip in 10 mM MgCl_2_, or 1 mM N-OH-Pip in 10 mM MgCl_2_. After 48 hr, the three treated lower leaves and three untreated upper leaves (leaf number 11-13) were harvested, pooled, respectively, and then frozen in liquid nitrogen. Frozen tissue was pulverized and divided into two aliquots: one for qRT-PCR and the other for metabolic profiling. Three biological repeats were performed per treatment in two independent experiments.

### RNA Isolation and Quantitative real-time PCR (qRT-PCR)

Total RNA was isolated from leaves using Trizol reagent (Invitrogen) according to the manufacturer’s instructions. Two µg RNA were used to synthesize cDNA by oligo dT and reverse transcriptase. For qRT-PCR, each cDNA sample was amplified with gene-specific primers (Table S2) using Green Taq DNA polymerase (GenScript) with EvaGreen Dye (Biotium) and the MJ Opticon 2 (Bio-Rad). *UBC21 (Ubiquitin-Conjugating Enzyme 21; At5g25760)* mRNA abundance was used to normalize the expression value in each sample. The comparative Ct method (2^-ΔΔCt^) was performed to determine the relative expression. The fold change of each value was normalized to the value of MgCl_2_ treated local leaves of WT.

### SAR Assay

SAR bacterial growth assays were performed as described (Navarova et al., 2012) with slight modification. Each plant pot was drenched with 10 mL water, 1 mM L-(-)- Pipecolinic acid (Pip) (Oakwood, CA) or 1 mM N-OH-Pip. After 24 hr, three lower leaves of each plant were infiltrated with 10 mM MgCl_2_ or a 5×10^6^ cfu/ml suspension of *Pst avrRpt2* in 10 mM MgCl_2_. Two days later, one upper leaf of each plant was inoculated with a 1×10^5^ cfu/mL suspension of *Psm* ES4326, and then plants were covered with a dome to maintain humidity. The titer of *Psm* ES4326 in the upper leaves was quantified at 3 dpi by homogenizing leaves discs in 1 mL 10 mM MgCl_2_, plating appropriate dilutions on NYGA medium with rifampicin (100 µg/mL), incubating plates at 28°C for 2 days, and then counting bacterial colonies. Three plants were used per condition and the experiment was repeated more than 3 times.

### Symptom Analysis and Metabolic Profiling of *Pst avrRpt2* Infected Leaves After Root Treatments With Chemicals

Col-0 WT and *fmo1* plants were treated with 10 mL of water, 1 mM of Pip, or 1 mM of N- OH-Pip by root application. 24 hr later, one leaf of each WT and *fmo1* plant was inoculated with a 5×10^6^ cfu/ml suspension of *Pst avrRpt2* in 10 mM MgCl_2_ and then plants were covered. Symptoms of infected leaves were photographed 48 hpi. Four biological replicates were tested. For metabolic profiling, plants were similarly treated with chemicals for 24 hr and then three random leaves of each plant were harvested (- infection condition). Three of the remaining leaves of each plant were infiltrated with a 5×10^6^ cfu/mL suspension of *Pst avrRpt2* in 10 mM MgCl_2_ and harvested 48 hpi (+ infection condition). Ion abundance of Pip, N-OGlc-Pip, m/z = 100.075, camalexin, and SA-Glc were quantified using LC-MS. Four biological replicates were tested. The experiment was repeated twice with similar results.

### Minimum Inhibitory Concentration (MIC) Assay

MIC assays were performed according to (Sledz, 2015) with slight modification. A single colony of *Psm* ES4326 or *Pst* DC3000 was incubated in LB broth, while a single colony of *Pst avrRpt2* was incubated in NYGA broth without antibiotics and shaken (220 rpm) at 28°C for 24 hr. The culture was diluted to OD_595_ = 0.1 with LB and then shaken at 28°C until mid-log phase (OD_595_ = 0.5). Cultures were diluted to OD_595_ = 0.001 (∼10^6^ cfu/mL) and aliquots were transferred to wells of a 96-well microplate. Chemicals (Pip, N-OH-Pip, and salicylic acid; concentration range 0 - 1 mM) were added to each culture plate and then plates were shaken at 28°C. OD_595_ measurements of each culture were recorded at 0, 12, 24, 36 hr post-chemical treatment. For *Psm* ES4326 or *Pst* DC3000 assays, four biological repeats were performed per experiment in three independent trials. For *Pst avrRpt2* assays, four biological repeats were performed per experiment in two independent trials.

### Construction of FMO1 Mutants

The open reading frame of *FMO1* lacking the stop codon was amplified from *Arabidopsis* Col-0 WT cDNA by PCR using *FMO1* specific primers (Table S2) and cloned into the pCR8/GW/TOPO vector (Life Technologies, Carlsbad, CA). Two alanine substitution mutants, *FMO1*(*G17A/G19A*) and *FMO1*(*G215A*), were generated using pCR8/GW/TOPO-FMO1 as template and *fmo1* mutant primers (Table S2). All constructs were confirmed by DNA sequence analysis. WT and mutant *FMO1* cDNAs were subcloned into pEAQ-HT-DEST3 (Peyret and Lomonossoff, 2013) to create C-terminal 6x His-tagged fusion proteins. Plasmids were introduced into *E. coli* DH5 alpha and *A. tumefaciens* C58C1 by heat shock transformation.

### Transient Expression in *N. benthamiana*

*Agrobacterium* strains harboring the pEAQ-gene constructs were grown on LB agar plates with the appropriate antibiotics. After 48 hr of growth, cells were removed from plates using an inoculation loop and resuspended in 1 mL LB. Cells were centrifuged at 4000 x *g* for 5 min, the supernatant was removed, and cells were resuspended in 1 mL *Agrobacterium* induction medium (10 mM MES buffer, 10 mM MgCl_2_, 150 µM acetosyringone, pH 5.7) and incubated at room temperature with shaking for 2 hr. Cells were then diluted to a final OD_600_ of 0.3 in induction medium. In tests with supplemented Pip, cells were diluted to a final OD_600_ of 0.3 in induction medium + 1 mM Pip. These solutions were then infiltrated into the underside of *N. benthamiana* leaves (3 leaves per plant) using a needleless 1 mL syringe. Plants were grown on a growth shelf with a 16- hr light/ 8-hr dark cycle for 28 hr or 48 hr prior to sample harvest for metabolic analysis and immunoblot. Total protein of each sample was extracted from two leaf discs (1 cm diameter per disc) by using urea buffer (8M urea, 15% β-mercaptoethanol, 3x Laemmli buffer). Proteins were separated by 12% SDS-PAGE analysis and transferred to a PVDF membrane and visualized by Ponceau S red staining before immunoblot analysis. FMO1-6x-His, FMO1(G17A/G19A)-6x-His, FMO1(G215A)-6x-His proteins were visualized by chemiluminescence using anti-His (Qiagen), peroxidase-conjugated secondary antibodies (Bio-Rad), and ECL reagent (GE Biosciences).

### Electrolyte Leakage and Hypersensitive Reaction Assays

Electrolyte leakage and hypersensitive reaction (HR) assays were performed according to (Cheong et al., 2014). 30-32 -day-old WT and *fmo1* plants were irrigated with 10 mL water, 1 mM Pip, or 1 mM N-OH-Pip. One day later, six to seven random leaves of each plant were inoculated with a 3×10^8^ cells/mL suspension of *Pst* DC3000 (vector) or *Pst* DC3000 (*avrRpt2*) and then incubated at room temperature under lights. For electrolyte leakage assay, at 5 hr post-inoculation, three leaf discs (7 mm diameter) of each plant were pooled and floated in 20 mL of water in petri dishes. 5 min later, the leaf discs were transferred to a 15 mL tube containing 3 mL water and incubated at room temperature for 1 hr with shaking. Conductivity of each sample before and after boiling was measured using an electrical conductivity meter (Spectrum Technologies). The percentage of electrolyte leakage was calculated as conductivity before boiling/conductivity after boiling. For HR assays, leaf phenotypes of WT and *fmo1* for each condition were record at 8 hpi. Three plants were used per condition and the experiment was repeated 3 times.

---

**Table E1.** Major Product Ions of Metabolites Measured in this Study.

| Metabolite | m/z | Adduct | Retention time<br>(min) | Technique |
| --- | --- | --- | --- | --- |
| N-OGlc-Pip | 308.134 | $[M+H]^+$ | 2.1 | Reverse-phase |
| N-OH-Pip | 146.081 | $[M+H]^+$ | 2.2 | Hydrophilic-interaction |
| Pip | 130.086 | $[M+H]^+$ | 3.8 | Hydrophilic-interaction |
| m/z =<br>100.075 | 100.075 | $[M+H]^+$ | 1.0 | Hydrophilic-interaction |
| Camalexin | 201.048 | $[M+H]^+$ | 16.4 | Reverse-phase |
| SA-Glc | 323.0742 | $[M+Na]^+$ | 5.5 | Reverse-phase |

